# Interstitial telomeric sequences promote gross chromosomal rearrangement via multiple mechanisms

**DOI:** 10.1101/2024.04.11.589032

**Authors:** Fernando R. Rosas Bringas, Ziqing Yin, Yue Yao, Jonathan Boudeman, Sandra Ollivaud, Michael Chang

**Affiliations:** European Research Institute for the Biology of Ageing, University of Groningen, University Medical Center Groningen, Groningen, the Netherlands

**Keywords:** interstitial telomeric sequence, gross chromosomal rearrangement, de novo telomere addition, telomeric DNA replication, DNA repair

## Abstract

Telomeric DNA sequences are difficult to replicate. Replication forks frequently pause or stall at telomeres, which can lead to telomere truncation and dysfunction. In addition to being at chromosome ends, telomere repeats are also present at internal locations within chromosomes, known as interstitial telomeric sequences (ITSs). These sequences are unstable and prone to triggering gross chromosomal rearrangements (GCRs). In this study, we quantitatively examined the effect of ITSs on GCR rate in *Saccharomyces cerevisiae* using a genetic assay. We find that GCR rate increases exponentially with ITS length. This increase can be attributed to the telomere repeat binding protein Rap1 impeding DNA replication and a bias of repairing DNA breaks at or distal to the ITS via de novo telomere addition. Additionally, we performed a genome-wide screen for genes that modulate the rate of ITS-induced GCRs. We find that mutation of core components of the DNA replication machinery leads to an increase in GCRs, but many mutants known to increase GCR rate in the absence of an ITS do not significantly affect GCR rate when an ITS is present. We also identified genes that promote the formation of ITS-induced GCRs, including genes with roles in telomere maintenance, nucleotide excision repair, and transcription. Our work thus uncovers multiple mechanisms by which an ITS promotes GCR.

**Significance statement:** Telomeric DNA repeats are found at the ends of linear chromosomes where they, together with specialized proteins that bind to them, protect chromosome ends from degradation and unwanted DNA repair activities. Telomeric repeats can also be found at internal locations in the genome, where they are called interstitial telomeric sequences (ITSs). ITSs are prone to breakage and are associated with human diseases. In this study, using baker’s yeast as a model organism, we show that instability at ITSs is driven by multiple factors, and identify genes that either promote or suppress gross chromosomal rearrangements induced by the presence of an ITS.

## Introduction

Telomeres consist of repetitive DNA sequences and associated proteins, and are essential structures that ‘cap’ and protect the ends of eukaryotic chromosomes (1). A dysfunctional telomere will cause the chromosome end to be recognized as DNA damage, activating the DNA damage response and inhibiting cell proliferation. Telomeres, however, are also problematic because they are difficult to replicate. First, due to the enzymatic characteristics of DNA polymerases, the standard DNA replication machinery cannot fully replicate DNA ends—a problem famously referred to as the end replication problem (2). This problem is solved by an enzyme called telomerase, which adds telomeric repeats to the ends of the chromosomes (3). A second (and less-appreciated) problem is that DNA replication forks often stall and collapse while traversing telomeric sequences. This is highly conserved throughout evolution, having been reported in budding yeast *Saccharomyces cerevisiae*, fission yeast *Schizosaccharomyces pombe*, and mouse and human cells (4–7).

There are at least three obstacles that could hinder the progression of the replication fork through telomeric sequences: 1) proteins that are tightly bound to telomeric sequences; 2) secondary structures (e.g. G-quadruplexes) that form due to the repetitive G-rich nature of telomeres (8); and 3) RNA-DNA hybrids formed from the transcription of telomeric repeat-containing RNA (TERRA; 9). If such impediments to replication fork progression are not properly dealt with, telomeres can become truncated and dysfunctional.

The difficulty of replicating telomeric repeats extends to telomeric sequences located at internal regions of the genome, called interstitial telomeric sequences (ITSs). ITSs are naturally found in the genomes of many species and are thought to be either remnants of ancestral chromosomal fusions or erroneous insertions of telomeric repeats that occur during the repair of DNA double-strand breaks (DSBs; 10, 11). The human genome contains over 80 ITSs that have at least four telomeric repeats (12). ITSs are especially hazardous to genome stability because they are prone to breakage, provoke genome rearrangements, and are often found at translocation breakpoints in cancer cells (11, 13, 14). A break at an ITS can also be ‘healed’ by telomerase, resulting in the generation of a new telomere and loss of genetic material distal to the ITS. Such de novo telomere addition events have been linked to human disorders, including terminal deletions of chromosome 16 (alpha-thalassemia; 15, 16) and chromosome 12 (mental retardation; 17).

In this study, we quantitatively measured ITS stability in *S. cerevisiae* using a gross chromosomal rearrangement (GCR) assay. We find that GCR rates rise exponentially with increasing ITS length. This increase is caused by the presence of the telomere repeat binding protein Rap1, which is known to impede DNA replication (5, 18–20), and a strong preference for repairing DNA breaks at or distal to the ITS by de novo telomere addition. In addition, we performed a genome-wide screen to identify genes that contribute to ITS stability. We find that mutations in DNA replication genes elevate GCR rates, but many genes reported as general GCR suppressors do not exhibit an effect in the presence of an ITS. We also identified genes that promote ITS-induced GCRs, including genes important for telomere maintenance, nucleotide excision repair, and transcription. Our work reveals the multiple mechanisms by which an ITS promotes GCR.

## Results

### GCR rate increases exponentially with ITS length

To quantitatively assess ITS stability, we modified the GCR assay developed by Chen and Kolodner by inserting an ITS of varying length between the most telomere-proximal essential gene on the left arm of chromosome V (*PCM1*) and two counterselectable markers (*CAN1* and *URA3*) used in the original assay (21; Figure 1A). The GCR rate (GCR events per cell division) can be determined by measuring the rate of simultaneous loss of *CAN1* and *URA3*—detected by growth on canavanine and 5-fluoroorotic acid (5-FOA), respectively— with a fluctuation test. We observe that the GCR rate increases with ITS length in two distinct exponential phases (Figure 1B; Table S1). Up to 50 bp, the GCR rate increases rapidly (1.6×10^-9^ with no ITS to 4.5×10^-6^ with a 50-bp ITS); beyond 50 bp, the GCR rate still increases exponentially, but more modestly (to 1.5×10^-4^ with a 300-bp ITS). To rule out the possibility that the increase in GCR rate is simply due to the high G/C-content of the ITS, we inserted 300 bp of lambda phage DNA, which has a 62% G/C content, similar to wild-type telomeric sequence. The GCR rate of this strain is similar to that of the ‘no-ITS’ control (Figure 1C; Table S1). Telomeric sequences promote transcriptional silencing of adjacent genes (22). To determine whether the increase in GCR rate was in part caused by silencing of the *CAN1* and *URA3* genes, we measured the GCR rate in a 300-bp ITS-containing strain deleted for *SIR2*, a gene needed for silencing (23). We find that absence of Sir2 does not affect the GCR rate (Figure 1C; Table S1). Thus, the ITS-induced increase in GCR rate is due neither to the high G/C-content of the ITS nor to silencing of the GCR markers.

**Figure 1.**
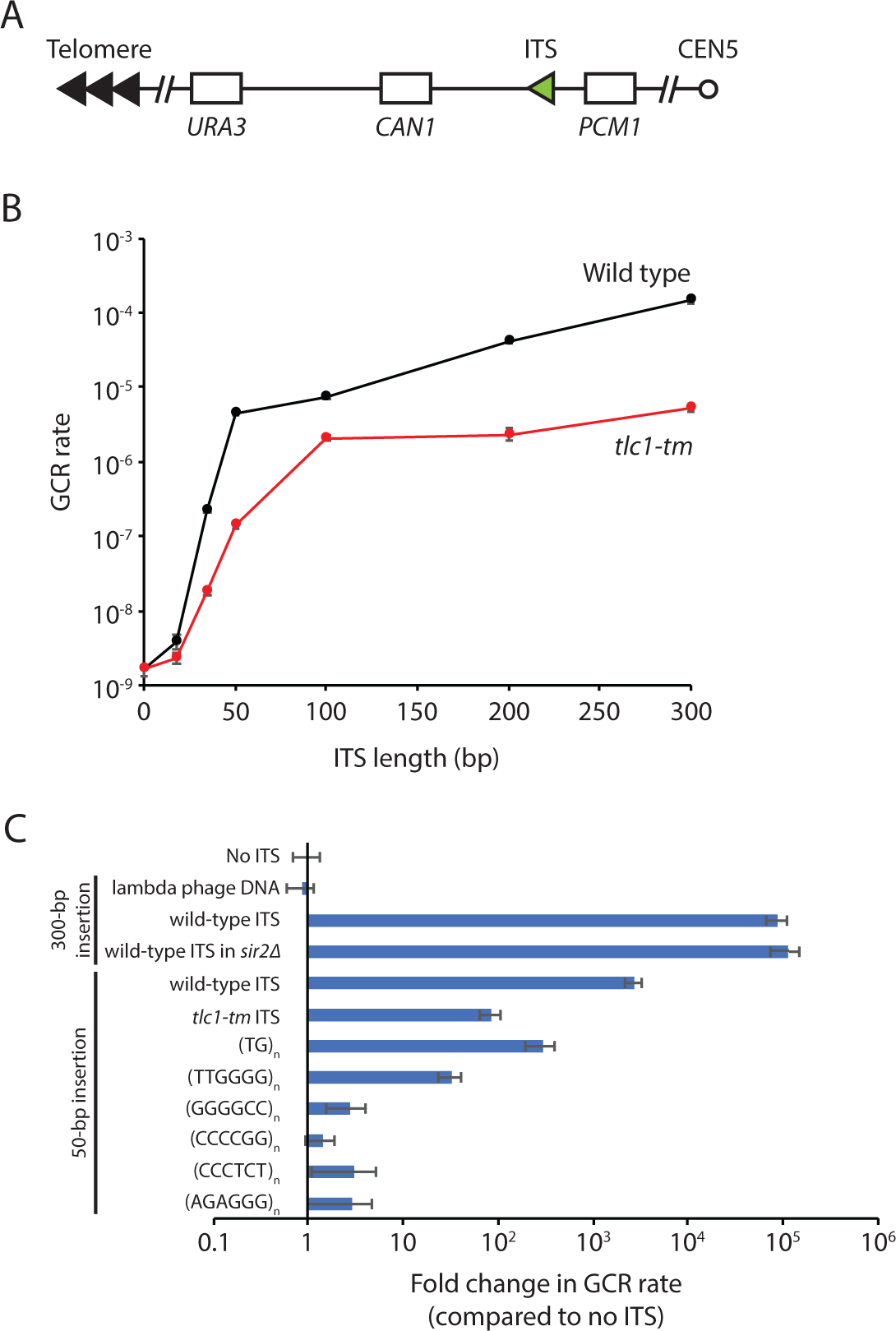
GCR rate increases with ITS length due to Rap1-dependent inhibition of replication and a bias of repairing DNA breaks via de novo telomere addition. (**A**) Schematic diagram of the ITS-GCR assay. Loss of both *URA3* and *CAN1* allows growth on 5-FOA and canavanine. ITSs of varying length were inserted between *CAN1* and *PCM1*, the most telomere-proximal essential gene on the left arm of chromosome V. (**B**) GCR rate plotted as a function of ITS length. Wild-type ITS in black; *tlc1-tm* ITS in red. (**C**) GCR rates of strains with the indicated sequence insertions are plotted. Error bars represent SEM (n = 3 or 4).

Two main factors can explain why GCR rate increases with ITS length. First, telomeric DNA is difficult to replicate, so the longer the ITS, the greater the possibility for a GCR event to occur. Second, if a DNA break occurs at or distal to the ITS, the probability that the break is ‘repaired’ by de novo telomere addition (resulting in a GCR) is very high if at least 34 bp of telomeric sequence is present on the centromere-side of the break. A sharp threshold separates a DSB from a telomere; DNA ends with 34 bp of telomeric sequence are recognized as critically short telomeres and efficiently extended by telomerase, whereas those with less than 34 bp are not (24). The bias towards repair via de novo telomere addition reaches a maximum once the ITS is sufficiently longer than this threshold; once a DNA end looks like a telomere, it cannot look even more like a telomere—at least in terms of de novo telomere addition. Thus, the increase in GCR rate is more modest once the ITS is longer than 50 bp.

Note that the inflection point does not occur at 34 bp because a break could occur either within the ITS or distal to the ITS. For a 34-bp ITS, the break that leads to the de novo telomere addition most likely occurs distal to the ITS; if it were to occur within the ITS, the resulting DNA end would have less than 34 bp of telomeric sequence and would therefore not be efficiently extended by telomerase. For increasingly longer ITSs, the probability that the break occurs within the ITS increases, as does the probability of that break leaving at least 34 bp of telomeric sequence on the centromere-side of the break. Both of these probabilities will eventually reach a maximum (i.e. at a certain ITS length, almost all breaks will occur within the ITS, and almost all of these breaks will leave at least 34 bp of telomeric sequence on the centromere-side of the break), and further increases in ITS length will not affect these factors (although GCR rate will continue to increase due to Rap1-mediated fork collapse; see below). We do not know the exact ITS length where this effect occurs, but the net effect is an inflection point in the GCR rate curve at approximately 50 bp in ITS length.

### Rap1 promotes the ITS-induced increase in GCR rate

The difficulty in replicating telomeric DNA has been attributed to the binding of Rap1 to telomeric repeats (5, 18–20). To test whether Rap1 is promoting the ITS-induced increase in GCR rate, we examined the effect of an ITS with *tlc1-tm* [(TG)_0-4_TGG]_n_ATTTGG telomeric repeats (25), rather than wild-type (TG)_0-6_TGGGTGTG(G)_0-1_ repeats (26). The *tlc1-tm* mutant repeats disrupt Rap1 association, but can still be efficiently recognized by telomerase and cap telomeres when located at chromosome ends (27). We find that the *tlc1-tm* ITS also increases GCR rate, but to a lesser extent than the wild-type ITS (Figure 1B; Table S1). The relationship between GCR rate and *tlc1-tm* ITS length is consistent with the DSB-telomere threshold, which is not affected by Rap1 binding (24), playing a dominant role. As *tlc1-tm* ITS length increases, repair by de novo telomere addition increases, reaching a maximum once the *tlc1-tm* ITS is sufficiently longer than the DSB-telomere threshold. However, the more modest exponential increase in GCR rate observed with wild-type ITS lengths above 50 bp is not seen with the *tlc1-tm* ITS. Since Rap1 is not completely depleted from *tlc1-tm* sequences (27), we also measured the effect of a 50-bp insertion of TG dinucleotides, i.e. (TG)_25_, which is predicted to disrupt Rap1 binding even more so than *tlc1-tm* repeats (28), but like *tlc1-tm*, can still be efficiently extended by telomerase (24). We find that the (TG)_25_-induced GCR rate is comparable to the 50-bp *tlc1-tm* ITS-induced GCR rate; i.e. there is an increase in the GCR rate, but much less than with a 50-bp wild-type ITS (Figure 1C; Table S2). The difficulty in replicating telomeric sequences has also been attributed to the formation of G-quadruplexes, which is compromised in TG dinucleotide and *tlc1-tm* repeats (29). Thus, we also tested 50 bp of TTGGGG *Tetrahymena* telomeric repeats, which can be extended by *S. cerevisiae* telomerase (30) and have the ability to form G-quadruplexes (31), but does not bind Rap1 (32). We find that this sequence increases GCR rate to a level slightly less than that of the 50-bp *tlc1-tm* ITS (Figure 1C; Table S2). These findings suggest that Rap1, rather than G-quadruplex formation, promotes the wild-type ITS-induced increase in GCR rate.

To further assess the ability of G-quadruplexes to induce GCR, we measured the effect of inserting 50 bp of the hexanucleotide repeats GGGGCC/GGCCCC and CCCTCT/AGAGGG on GCR rate. Expansion of these repeats cause neurogenerative diseases (amyotrophic lateral sclerosis and frontotemporal degeneration for GGGGCC/GGCCCC and X-linked dystonia parkinsonism for CCCTCT/AGAGGG), and the formation of G-quadruplexes by these repeats has been proposed to be involved in their expansion (33). The insertion of these sequences only mildly increases GCR rate (Figure 1C; Table S2). Our findings indicate that G-quadruplexes play a relatively minor role, in comparison to Rap1, in hindering the replication of *S. cerevisiae* telomeric DNA, in agreement with previous work, including an in vitro study examining the replication of telomeric sequences with and without Rap1 (5, 20).

### Breaks occur within and distal to the ITS

Since most GCR events detected in this assay without an ITS are predominantly de novo telomere addition events (34), we reasoned that the addition of an ITS would only further increase this bias, most likely with the new telomere added at the ITS. Thus, to gain insight into the mechanisms that underlie the increased GCR rates induced by the presence of an ITS, we sequenced the ITS region from independently isolated canavanine- and 5-FOA-resistant mutants (i.e. GCR survivors) derived from strains that either had the 50-bp ITS or the 100-bp ITS, 20 from each strain. As we expected, de novo telomere addition occurred at the ITS in all 40 GCR survivors. Since yeast telomerase adds imperfect, degenerate repeats (26), we can determine the exact position were telomerase added the new telomere by comparing with the original 50-bp and 100-bp ITS sequences (Figure 2A). Only a centromere-proximal portion of the ITS was retained in some of the GCR survivors (nine of 20 derived from the 50-bp ITS strain, 18 of 20 derived from the 100-bp ITS strain), suggesting that the DNA replication fork collapsed while traversing the ITS, followed by the telomerase-mediated addition of a new telomere. In all GCR survivors, at least 30 bp of the original ITS sequence was retained, consistent with DNA ends with less than 30 bp of telomeric sequence being recognized as DSBs and not efficiently extended by telomerase (24).

**Figure 2.**
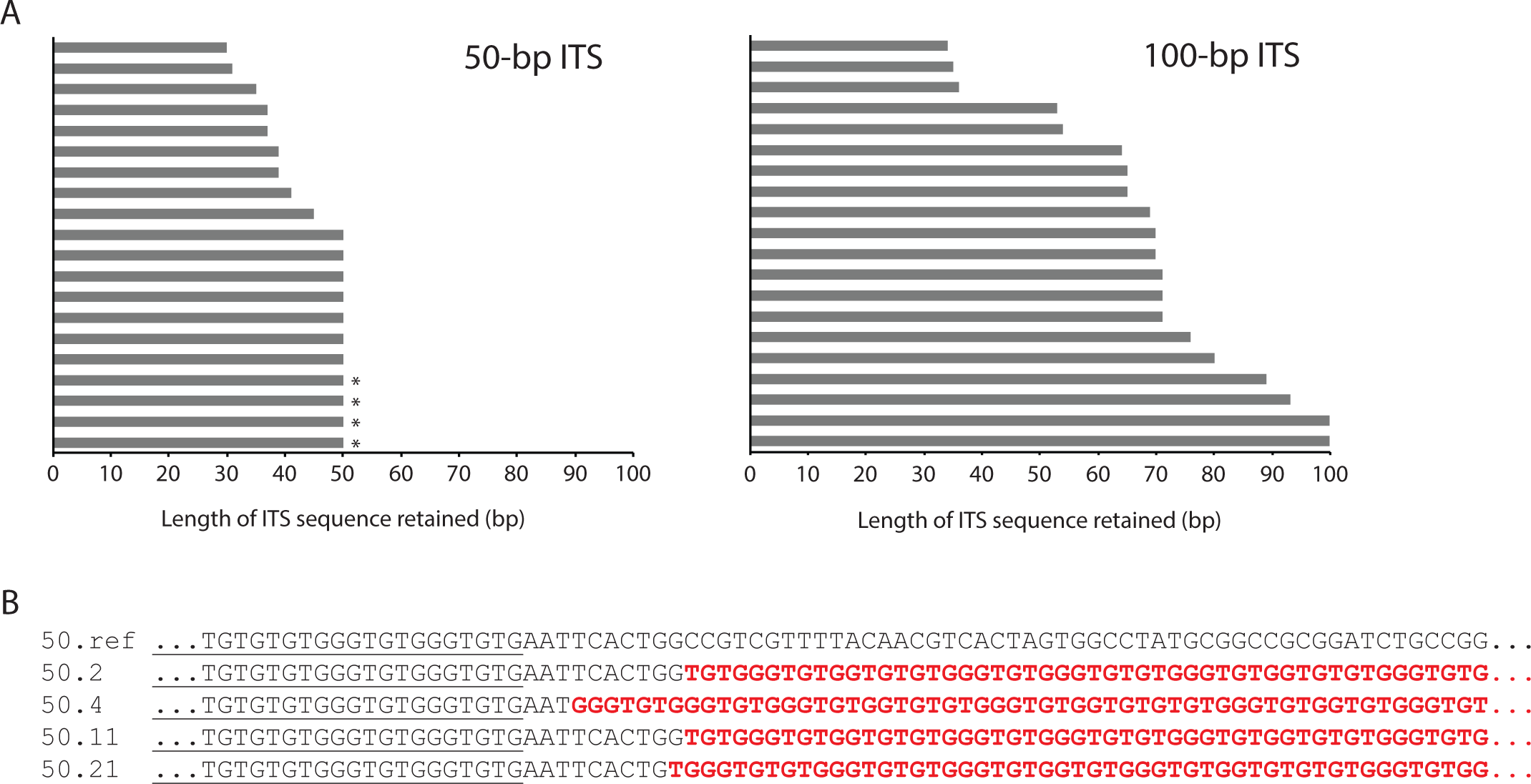
Breaks occur within and distal to the ITS. (**A**) GCR survivors were isolated by growing strains with either a 50-bp ITS or 100-bp ITS on agar plates containing canavanine and 5-FOA. Twenty independent isolates from each strain were analyzed. In all 40 isolates, the presence of a de novo telomere added at the ITS site was confirmed by PCR and sequencing. The length of the original ITS retained for each de novo telomere is plotted. Four isolates also retained sequence downstream of the ITS and are indicated by an asterisk. (**B**) The sequences of the four isolates are aligned to DNA sequence from the original 50-bp ITS-containing strain (50.ref). The last 20 bp of the ITS are underlined. Newly added telomeric sequence in the GCR survivors is in bold and red.

Surprisingly, the entire ITS sequence was retained in the remaining GCR survivors (11 of 20 derived from the 50-bp ITS strain, two of 20 derived from the 100-bp ITS strain). These findings suggest that the DNA replication fork likely collapsed distal to the ITS, followed by resection back to the ITS prior to telomere addition, leaving the original ITS intact. Consistent with our observations, it has been previously shown that sites of repair-associated telomere addition (SiRTAs; hot spots in the genome for de novo telomere addition) can be located several kilobases away from an induced DSB (35). In four GCR survivors derived from the 50-bp ITS strain, up to 10 bp immediately downstream of the ITS was also retained (Figure 2B). This is analogous to SiRTAs, which have a bipartite structure consisting of a Stim sequence and a Core sequence; Cdc13 can bind to a Stim sequence (in this case, the ITS) and stimulate de novo telomere addition at the downstream Core sequence (35).

### Identification of genes that suppress ITS-induced GCR events

To identify genes that suppress ITS instability, we used a high-throughput replica-pinning approach that we recently developed to detect low-frequency events (36), such as GCRs. First, we introduced the 50-bp ITS GCR assay components into the yeast knockout (YKO) and conditional temperature-sensitive (ts) strain libraries (37, 38) using the synthetic genetic array (SGA) method (39). The resulting strains were amplified by high-throughput replica-pinning, yielding 24 colonies per YKO or ts strain, onto media containing canavanine and 5-FOA to select for GCR events (see Materials and Methods for details). GCR frequencies (percentage of GCR-positive colonies) were calculated for each mutant strain (Figure 3A).

**Figure 3.**
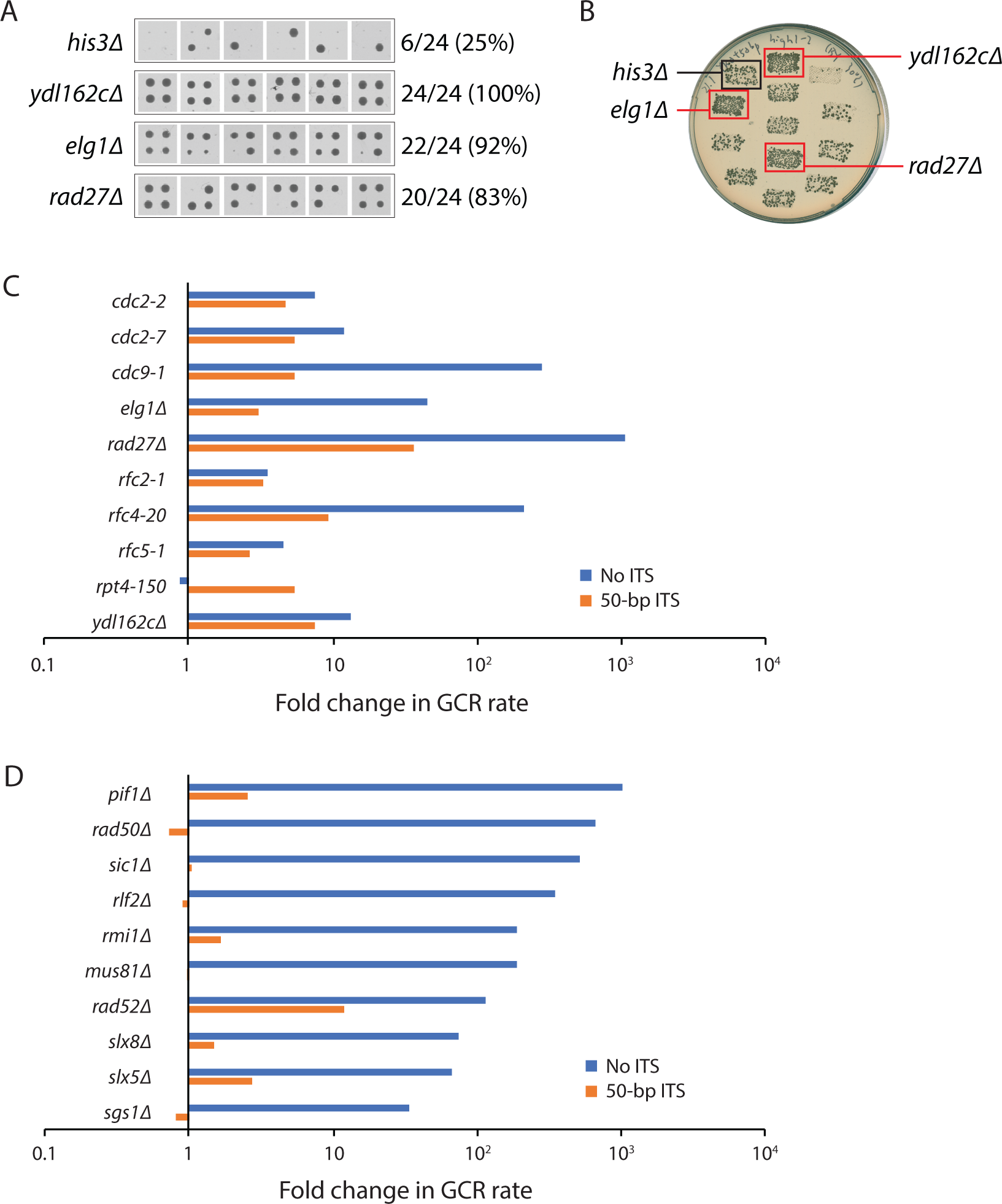
Identification of genes that suppress ITS-induced GCR rate. (**A**) A high-throughput screen was performed as described in the text. All 24 replica-pinned colonies on media containing both canavanine and 5-FOA of the *his3Δ* control strain and three selected mutants with increased GCR frequencies are shown. (**B**) Putative hits from the high-throughput screen were tested in a patch-and-replica-plate assay. An example plate, with three mutants that tested positive (red boxes), is shown. A negative control (*his3Δ*) and a positive control (*ydl162cΔ*) were included on each plate. (**C**) Fold change in GCR rate of mutants identified in the ITS-GCR screen. (**D**) Fold change in GCR rate of ten known GCR suppressors not identified in the ITS-GCR screen. Fold change of mutants without an ITS is relative to a wild-type strain without an ITS. Fold change of mutants with a 50-bp ITS is relative to a wild-type strain with an ITS. Data for YKO strains without an ITS were taken from a previous publication (50).

The median GCR frequency was 38% for the YKO strains and 42% for the ts strains. Only mutants with a GCR frequency above 70% (corresponding to a Fisher’s exact test P-value of <0.002) were analyzed further (Dataset S1). A total of 54 YKO and 44 ts mutants met this criterion. These strains were generated again by SGA and further tested using a patch-and-replica-plate approach (Figure 3B; 40), from which 21 mutants (5 YKO and 16 ts) appeared to have an increased ITS-induced GCR rate. Their GCR rates were determined by fluctuation tests. Ten of the 21 mutants have more than a twofold increase in ITS-induced GCR rate (Figure 3C; Table S3).

Of these nine genes (two of the mutants are different alleles of *CDC2*), eight are primarily involved in DNA replication (*CDC2*/*POL3*, *CDC9*, *ELG1*, *RAD27*, *RFC2*, *RFC4*, *RFC5*, and *YDL162c*). Pol3 is the catalytic subunit of DNA polymerase δ (41, 42). Rfc2, Rfc4, and Rfc5 are small subunits of the replication factor C (RFC) complex, which catalyzes the loading of the proliferating cell nuclear antigen (PCNA) sliding clamp (43). Elg1 forms an alternative RFC complex with Rfc2–5 that is implicated in unloading PCNA (44). Rad27 is a flap endonuclease involved in Okazaki fragment processing and maturation (45). Cdc9 is DNA ligase I, responsible for ligating Okazaki fragments together (46). *YDL162c* is an open reading frame that overlaps with the *CDC9* promoter; its deletion reduces *CDC9* expression (47). The ninth gene, *RPT4*, encodes an ATPase of the 19S regulatory particle of the 26S proteasome (48), which is important for many cellular functions, including DNA replication and genome stability (49).

With the exception of *RPT4*, all of these genes also suppress GCR rate in the absence of an ITS (Figure 3C; Table S3), which is not surprising given that these genes are involved in DNA replication; defects in DNA replication should increase GCRs regardless of whether an ITS is present. The ITS-specific increase in GCR rate observed in the *rpt4-150* strain may indicate that the proteasome removes a GCR-inducing protein that is bound to the ITS. An obvious candidate for such a protein is Rap1, although we do not detect a change in cellular Rap1 levels in *rpt4-150* cells (Figure S1). However, other explanations for the ITS-specific effect of *rpt4-150* are possible, and further studies are necessary to elucidate the mechanism.

### Many mutants increase GCR rate in the absence of an ITS but not in its presence

Many mutants have been reported to increase GCR rate in the absence of an ITS (50, 51). Interestingly, the majority of these mutants were not identified in our screen and might be false negatives. Therefore, we measured the GCR rate of 10 of these mutants in the presence of a 50-bp ITS (Figure 3D; Table S4). Seven of these mutants do not significantly alter ITS-induced GCR rate (i.e. less than 1.7-fold; *rad50Δ*, *sic1Δ*, *rlf2Δ*, *rmi1Δ*, *mus81Δ*, *slx8Δ, sgs1Δ*). For the remaining three, *pif1Δ*, *rad52Δ*, and *slx5Δ* increase ITS-induced GCR rate by 2.5-, 12-, and 2.7-fold, respectively, far lower than their effect on GCR rate in the absence of an ITS. Thus, it appears that many known GCR suppressors have a greatly reduced, if not completely absent, role in suppressing GCRs when an ITS is present. We also tested *rrm3Δ* because Rrm3 is a helicase that promotes replication fork progression through telomeric sequences (4). Deletion of *RRM3* has been reported to exhibit a mild fourfold increase in GCR rate in the absence of an ITS (52); we find that this is also true in the presence of an ITS (Table S4).

Most known GCR suppressors have roles in DNA replication or repair. However, one exception that stands out is the cyclin-dependent kinase inhibitor Sic1. Cells lacking Sic1 initiate DNA replication from fewer origins of replication, causing an increase in the distance a replication fork has to travel and in the time needed to replicate the entire genome (53), including the region of chromosome V relevant for the GCR assay (54). It was suggested that ongoing replication as *sic1Δ* cells enter anaphase may lead to an increase in DSBs and GCRs (53). ITS-induced pausing of the replication fork would also cause a delay in replicating DNA distal to the ITS, resulting in under-replication and DNA breaks distal to the ITS, consistent with the data we obtained by sequencing GCR survivors (Figure 2). The lack of an increase in GCR rate in *sic1Δ* strains with a 50-bp ITS suggests that the ITS is epistatic to *sic1Δ* in terms of delaying replication of the left arm of chromosome V. In other words, replication of the chromosomal region distal to the ITS is chronically delayed, so much so that further deletion of *SIC1* cannot make it worse.

### The *rfc2-1* and *rfc5-1* mutations dramatically accelerate senescence in the absence of telomerase

Mutations that increase ITS-induced GCR rate likely increase the probability of replication fork collapse within the ITS. Such mutations may also increase fork collapse at native telomeres, which could lead to truncation of telomeres and accelerated senescence in the absence of telomerase. Consistent with this idea, *cdc2-2* and *rad27Δ* have both been reported to cause accelerated senescence in the absence of telomerase (55, 56). We tested the remaining eight mutations that increase ITS-induced GCR rate. We sporulated diploids that are heterozygous in one of the genes of interest as well as *EST2*, which encodes the protein catalytic subunit of telomerase (57), and followed the growth of the haploid meiotic progeny by serial propagation in liquid cultures for several days (Figure 4). As expected, *est2Δ* cultures grew slower as the experiment progressed and cells senesced, but growth was eventually restored upon the emergence of survivors that utilize recombination-mediated mechanisms to maintain telomeres (58). We find that *est2Δ rfc2-1* and *est2Δ rfc5-1* double mutants senesce much faster than *est2Δ* single mutants. The *cdc2-7*, *cdc9-1*, and *elg1Δ* mutations appear to modestly accelerate senescence, while the *rpt4-150* mutation causes a slight delay in senescence. Thus, seven of the 10 mutants found to elevate ITS-induced GCR rate also cause accelerated senescence in the absence of telomerase.

**Figure 4.**
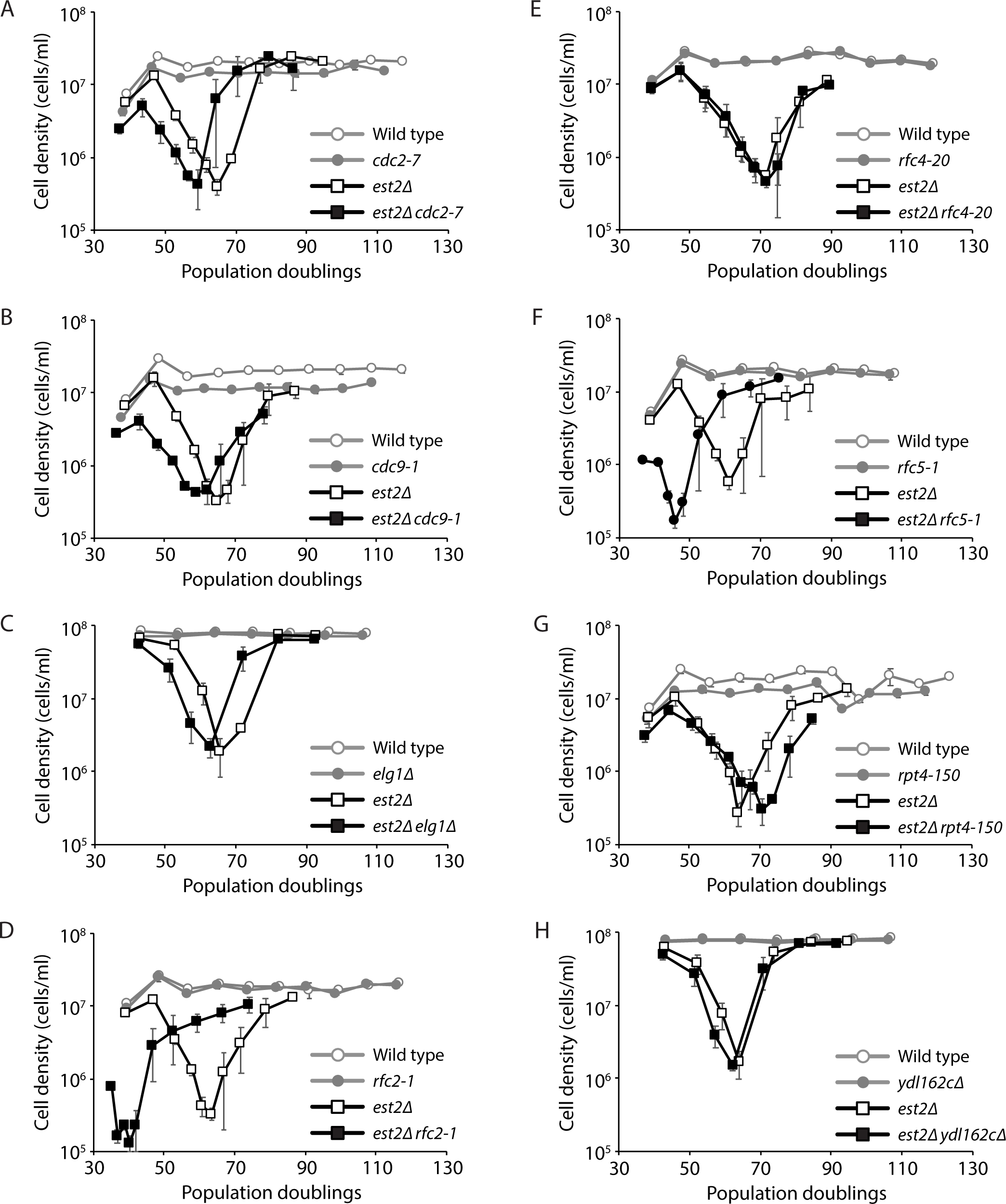
The *cdc2-7*, *cdc9-1*, *elg1Δ*, *rfc2-1*, and *rfc5-1* mutations accelerate senescence in the absence of telomerase. Senescence was monitored by serial passaging of haploid meiotic progeny derived from the sporulation of ZYY238 (**A**), ZYY236 (**B**), ZYY232 (**C**), ZYY244 (**D**), ZYY246 (**E**), ZYY248 (**F**), ZYY240 (**G**), and ZYY228 (**H**). Average cell density ±SEM of three independent isolates per genotype is plotted.

### Identification of mutants that decrease ITS-induced GCR rate

We repeated the genome-wide screen with minor adjustments that allowed us to identify mutants that decrease ITS-induced GCR rate (see Materials and Methods for details). The median GCR frequency was 79% for the YKO strains and 67% for the ts strains. Only mutants with a GCR frequency below 38% for the YKO strains and 31% for the ts strains (corresponding to a Fisher’s exact test P-value of <10^-4^) were analyzed further (Figure 5A; Dataset S2). A total of 248 YKO and 114 ts mutants met this criterion. These strains were generated again by SGA and further tested using the patch-and-replica-plate approach (Figure 5B). A large number of the remaining genes have roles in chromosome segregation, spindle assembly checkpoint, and microtubule-based processes. The identification of these genes is due to the *CIN8* gene located distal to the ITS on chromosome V (Yao and Yin et al., accompanying manuscript). De novo telomere addition at the ITS results in the loss of *CIN8*, which is synthetic lethal with mutation of these genes. Thus, we eliminated the hits that are known to be synthetic lethal with *cin8Δ*. We also re-tested these mutants with the patch-and-replicate-plate approach after introducing an extra copy of *CIN8* located elsewhere in the genome, and eliminated hits that did not show a decrease in ITS-induced GCRs. Finally, we performed an SGA analysis of the hits, identifying and eliminating those that showed a negative synthetic genetic interaction with truncation of chromosome V at the ITS. The remaining 180 (122 YKO and 58 ts) hits show a high degree of enrichment for genes involved in nucleotide excision repair and transcription (Figure 5C; Table S5). A subset of the mutants was subjected to fluctuation tests to determine their ITS-induced GCR rates (Figure 5D; Tables S5 and S6).

**Figure 5.**
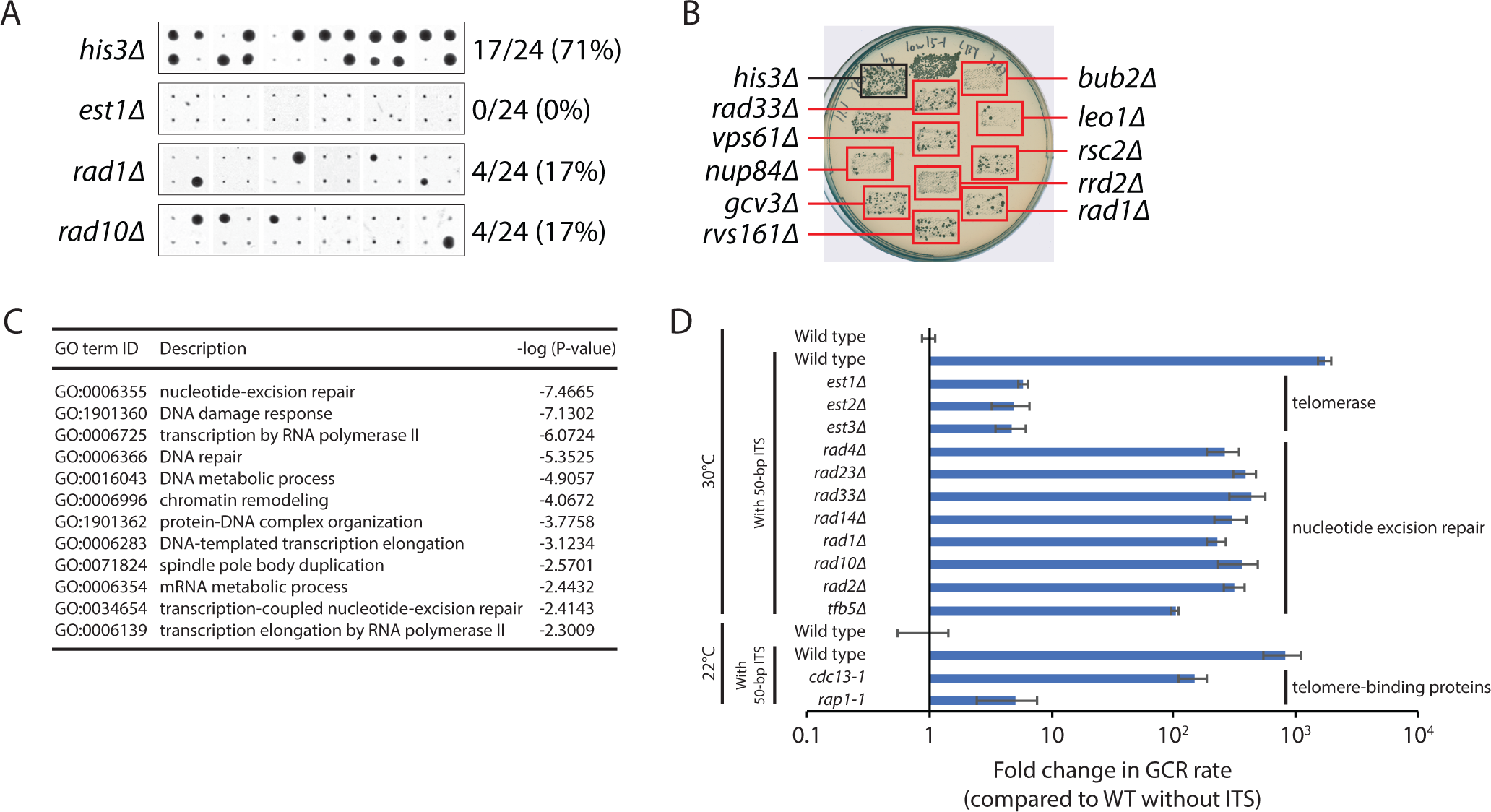
Identification of genes that promote ITS-induced GCR rate. (**A**) A high-throughput screen was performed as described in the text. All 24 replica-pinned colonies on media containing both canavanine and 5-FOA of the *his3Δ* control strain and three selected mutants with decreased GCR frequencies are shown. (**B**) Putative hits from the high-throughput screen were tested in a patch-and-replica-plate assay. An example plate, with ten mutants that tested positive (red boxes), is shown. A negative control (*his3Δ*) and a positive control (*bub2Δ*) were included on each plate. (**C**) The statistically supported GO terms enriched in the genes that promote ITS-induced GCR rate are shown. (**D**) GCR rates of the indicated strains containing a 50-bp ITS are plotted. Error bars represent SEM (n = 3–6).

As expected, our screen identified *EST1*, *EST2*, and *EST3*, which encode the three protein subunits of telomerase (59). Deleting either *EST1*, *EST2*, or *EST3* has no significant effect on GCR rate in the absence of an ITS (60). In contrast, in the presence of an ITS, deletion of *EST1*, *EST2*, or *EST3* decreases GCR rate by approximately 300-fold. However, *est1Δ*, *est2Δ*, and *est3Δ* strains with a 50-bp ITS have GCR rates that are still fivefold higher than a wild-type strain without an ITS, indicating that a 50-bp ITS can induce the formation of GCRs that are not the result of telomerase-mediated de novo telomere addition. These GCRs are still likely triggered by Rap1-mediated inhibition of replication, causing breaks either within or distal to the ITS that are repaired in a manner that leads to other types of GCRs, such as nonreciprocal translocations or interstitial deletions.

The *cdc13-1* and *rap1-1* mutants exhibit 5.5- and 159-fold decreases, respectively, in GCR rate in the presence of a 50-bp ITS. The *rap1-1* allele encodes a Rap1 mutant protein that is compromised in binding DNA (61), and may therefore be less capable of obstructing DNA replication and inducing GCR. The *cdc13-1* result is consistent with our previous observation that this mutant is defective in generating a new telomere at a DNA end with 34 bp of telomeric sequence (24). Cdc13 binds to single-stranded telomeric DNA and is important for preventing excessive resection of telomere ends and for telomerase recruitment (62, 63). The *cdc13-1* allele is defective in the former function (62). It has been reported that defective resection promotes de novo telomere addition (64), so it is possible that excessive resection in the *cdc13-1* mutant impairs de novo telomere addition.

We also identified many genes (*RAD1*, *RAD2*, *RAD4*, *RAD10*, *RAD14*, *RAD23*, *RAD33*, and *TFB5*) required for nucleotide excision repair (NER). Deletion of *RAD1* and *RAD10* has previously been reported to reduce GCR formation in mutant strains with elevated GCR rates (65). Rad1 and Rad10 form a structure-dependent endonuclease that cleaves 3’ flaps at junctions between single- and double-strand DNA during NER as well as homology-dependent repair processes (66). It was proposed that Rad1-Rad10 may cleave 3’ flaps during the formation of a GCR, including the addition of a de novo telomere where the initiating break occurs distal to the addition site (65; Figure S2A). If this model is correct, the fraction of GCR survivors that retain the entire ITS should decrease in *rad1Δ* or *rad10Δ* mutants. However, we find that this is not the case (Figure S2B), arguing against this model. Furthermore, the identification of many other genes involved in NER, which are not necessary for cleavage of 3’ flap structures, suggests instead that Rad1-Rad10 promote GCR formation within the context of the NER pathway.

NER can be divided into two subpathways: global-genome NER (GG-NER) and transcription-coupled NER (TC-NER). The NER genes identified in the screen are important for both subpathways. In GG-NER, Rad7 and Rad16 form a stable complex that is needed for transcription-independent recognition of UV-induced damage, while Rad26 is involved in TC-NER (67). Deletion of *RAD7* or *RAD16* has no effect on GCR rate, while deletion of *RAD26* decreases GCR rate approximately threefold (Table S6), suggesting the involvement of TC-NER. It has been reported that R-loops, consisting of an RNA-DNA hybrid and displaced single-stranded DNA that can arise during transcription, are processed into DSBs by TC-NER (68). The identification of many genes with functions in transcription in our screen is consistent with R-loops promoting GCR via TC-NER. Regardless of the precise mechanism, the role of TC-NER in promoting GCR is independent of the ITS, because deletion of *RAD1* or *RAD10* also reduces GCR formation in strains without an ITS present (65).

## Discussion

In this study, we examined ITS stability using a GCR assay in *S. cerevisiae*. We find that GCR rate increases exponentially with ITS length, which can be attributed to Rap1-mediated inhibition of replication of telomeric sequence, and to a bias in repairing DNA breaks by de novo telomere addition. The latter effect maximizes once the ITS is approximately 50 bp in length, as the ITS becomes longer than the DSB-telomere transition length of 34 bp. In addition, we performed a genome-wide screen and identified nine genes that contribute to the stability of telomeric sequences, most of whom encode core components of the DNA replication machinery. Surprisingly, many genes previously reported to suppress GCRs in the absence of an ITS do not do so in the presence of an ITS, highlighting the ability of an ITS to override mechanisms that normally suppress GCRs. We also find that genes with roles in telomere maintenance, NER, and transcription can facilitate the formation of ITS-induced GCRs.

### Binding of Rap1 to an ITS promotes its instability

Our findings confirm and extend previous studies showing that DNA-bound Rap1, much more so than G-quadruplex formation, hinders replication fork progression through telomeric sequences and induces DNA breakage (5, 18, 20). We show that these breaks can occur within the ITS, but also distal to the ITS. The breaks distal to the ITS are likely caused by incomplete replication of DNA distal to the ITS because of Rap1-mediated delay of fork progression at the ITS. Unexpectedly, we find that the GCR rate increases exponentially with ITS length (Figure 1B). This suggests that Rap1 molecules are acting in a cooperative manner to obstruct DNA replication. The Rap1-interacting proteins Rif1 and Rif2 connect neighboring and distal DNA-bound Rap1 units to form a high-order interlinked scaffold (69), the properties of which could change as telomere length increases. However, DNA-bound Rap1 obstructs replication independently of Rif1 and Rif2 (5, 18, 20). Rap1 has been reported to interact with DNA in multiple binding modes (70) and can, by itself, induce local DNA stiffening (71). Further studies are needed to determine whether these properties of Rap1 can explain why GCR rate increases exponentially with telomere length.

### Defects in DNA replication increase ITS-induced GCR rate and accelerate replicative senescence

We identified nine genes that suppress ITS-induced GCR rate (Figure 3C). Most have a direct function during DNA replication. If these mutants increase GCR rate by increasing DNA breakage, they may also increase breaks at native telomeres that would accelerate senescence in the absence of telomerase. This is already known to be true for *cdc2-2* and *rad27Δ* (55, 56), and we show that *cdc2-7*, *cdc9-1*, *elg1Δ*, *rfc2-1*, and *rfc5-1* also do so, especially the latter two (Figure 4). The dramatic effect of *rfc2-1* and *rfc5-1* on senescence is somewhat surprising given that they do not have the largest effects on GCR rate, with or without an ITS present, and the *rfc4-20* mutant does not significantly affect senescence at all. However, analysis of the genetic interaction profile similarity subnetwork for the identified genes, extracted from the global network (72), reveals that *rfc2-1* and *rfc5-1*, while clustering among DNA replication and repair genes, are positioned further away than the other identified replication genes (Figure S3), indicating related but distinct functions. Consistent with this idea, Rfc2 and Rfc5 are unique among the small RFC subunits in that they are about twice as abundant as Rfc3 and Rfc4 (73), and can form a stable heterodimer that acts on PCNA in vitro (74). *rfc2-1 rfc5-1* double mutants are inviable, and overexpression of *RFC5* can suppress the temperature sensitivity of *rfc2-1* (75). Furthermore, the temperature sensitivity and replication defect of the *rfc5-1* mutant can be suppressed by overexpression *POL30*, which encodes PCNA (76), suggesting that there is a deficiency in PCNA loading in this mutant. Thus, insufficient loading of PCNA may lead to replication defects that induce GCRs as well as accelerated senescence in the absence of telomerase.

It has been reported that the large subunit of RFC is aberrantly cleaved in Hutchinson-Gilford progeria syndrome (HGPS), resulting in defective loading of PCNA (77). HGPS is a disease characterized by accelerated aging and is caused by a mutation of the *LMNA* gene, leading to the expression of a truncated form of lamin A called progerin (78). At the cellular level, expression of progerin alters nuclear morphology and function, nucleolar activity, mitochondrial function, nucleocytoplasmic trafficking, and telomere homeostasis (79). It is still unclear how these alterations ultimately lead to the clinical features of HGPS, but telomere dysfunction may play a key role given that HGPS is associated with accelerated telomere erosion (80), and expression of telomerase improves the fitness of HGPS cells (81–83). Our findings support the hypothesis that premature senescence in HGPS is caused by a deficiency in PCNA loading and replication fork collapse at telomeres.

Interestingly, many of the other DNA replication genes identified in our screen have roles in lagging strand synthesis and Okazaki fragment processing. Elg1, in complex with Rfc2–5, unloads PCNA following Okazaki fragment ligation by Cdc9; depletion of Elg1 or Cdc9 causes accumulation of PCNA on DNA, which leads to genome instability (44, 84–87). Cells lacking Rad27 accumulate unligated Okazaki fragments (88), and presumably PCNA on DNA. Defects in lagging strand synthesis in the DNA polymerase δ mutants *cdc2-2* and *cdc2-7* would likely also increase the abundance of unligated Okazaki fragments and DNA-bound PCNA. Thus, a defect in unloading PCNA from DNA may also increase the probability of GCR.

### Presence of an ITS heavily biases repair towards de novo telomere addition

Surprisingly, our screen identified neither *PIF1* nor *RRM3*, which encode Pif1-family helicases. Stalling of replication forks traversing telomeric sequences increases in the absence of Rrm3, which is important for replication past nonhistone protein-DNA complexes (4, 89). Pif1 performs several functions that impact telomere biology, including inhibition of telomerase (90, 91), Okazaki fragment processing (92), unwinding of G-quadruplexes (93, 94), and promoting replication of Rap1-bound telomeric DNA (19, 20). Other genes encoding proteins known to suppress GCR in the absence of an ITS were also not identified, such as the MRX (Mre11-Rad50-Xrs2) complex, the STR (Sgs1-Top3-Rmi1) complex, the Mus81 endonuclease, and the SUMO-targeted ubiquitin ligase Slx5-Slx8. As they may have been false negatives in our screen, we tested deletions of some of these genes directly in the ITS-GCR assay.

Deletion of *PIF1*, which has been reported to cause an approximately 1000-fold increase in GCR rate in the absence of an ITS (60), only led to a 2.5-fold increase in the presence of a 50-bp ITS (Figure 3D; Table S4). We speculate that the role of Pif1 in suppressing GCR is primarily due to its function in suppressing de novo telomere addition (90), a function that is negated if the DNA end contains more than 34 bp of telomeric sequence (24). The residual 2.5-fold increase in the presence of the ITS could be due to the role of Pif1 in overcoming Rap1-dependent inhibition of DNA replication (19, 20), or its role in unwinding G-quadruplexes (93, 94). Consistent with that latter hypothesis, it has been previously reported that the GCR rate of the *pif1-m2* mutant, which is deficient in nuclear Pif1, is increased threefold in the presence of G-quadruplex motif sequences (94).

Strains lacking any member of the MRX complex have an approximately 600-fold increase in GCR rate in the absence of an ITS (21). Deletion of *SGS1* or *TOP3* increases GCR rate by 20–30-fold, while deletion of *RMI1* increases GCR rate by more than 170-fold (50, 95, 96). Deletion of *MUS81* increases GCR rate 100–200-fold, and deletion of *SLX5* or *SLX8* increases GCR rate by more than 60-fold (50, 65, 97). We find that deletion of either *RAD50*, *SGS1*, *RMI1*, *MUS81*, *SLX5*, or *SLX8* has a small or negligible effect on GCR rate in the presence of a 50-bp ITS (Figure 3D; Table S4). The MRX and STR complexes are central players in the repair of DSBs (98, 99); Mus81 is a structure-specific endonuclease that resolves replication and recombination intermediates (100); Slx5-Slx8 plays diverse roles in genome maintenance (101). In the absence of an ITS, disruption of these complexes will increase the likelihood of inappropriate repair that leads to a GCR. This effect is likely negated when the DNA break occurs at or distally adjacent to an ITS, since these breaks would be healed by de novo telomere addition regardless of whether these complexes are present. Consistent with this idea, a DSB that has telomeric repeats on one side of the DSB will result in asymmetric processing of the ends: the non-telomeric end will recruit the MRX complex and move to the nuclear pore complex while the telomeric side will not, favoring elongation by telomerase (102).

In summary, we have shown that ITSs are unstable and promote the formation of GCRs via multiple mechanisms. First, Rap1 impedes ITS replication, which increases the probability of fork collapse within the ITS. Second, impeding replication at the ITS will also increase fork collapse distal to the ITS due to incomplete replication of downstream DNA. Third, a DSB within or distal to the ITS, whether caused by fork collapse or other means, will bias repair towards de novo telomere addition, especially if at least 34 bp of telomere sequence remains on one side of the DSB. Lastly, transcription and NER can facilitate the formation of ITS-induced GCRs. Some of these mechanisms are also at play at native telomeres. For example, Rap1 also impedes replication at native telomeres, and a DSB within a telomere will be preferentially healed by telomerase, but this is not a problem precisely because it is desirable for a truncated telomere to be extended by telomerase. Thus, mechanisms that have evolved for proper telomere function are sources of instability and GCR at an ITS.

## Materials and methods

### Yeast strains and media

All yeast strains used in this study are listed in Table S7. To prevent the onset of replicative senescence, the *est* strains used in Figure 5C were generated by replacing the corresponding *EST* gene with a kanMX cassette in ZYY141, and then used immediately for fluctuation tests without storing the strains. Standard yeast genetic and molecular methods were used (103, 104). Experiments involving a strain containing a ts allele were performed at 22°C; in all other experiments, yeast strains were cultured at 30°C. ITS and other DNA sequences were inserted in GCR assay-containing strains at the *PRB1* locus and are listed in Table S8.

### Fluctuation tests of GCR rates

Fluctuation tests for the quantification of GCR rates were performed essentially as previously described (105) by transferring entire single colonies from YPD plates to 4 ml of YPD liquid medium. Cultures were grown to saturation (30°C for YKO strains and 22°C for ts mutants). 50 µl of a 10^5^-fold dilution were plated in YPD plates. An strain-dependent quantity of cells was plated on SD-arg+canavanine+5-FOA. Colonies were counted after incubation at 30ᵒC or 22ᵒC for 3–7 days. The number of GCR (canavanine- and 5-FOA-resistant) colonies was used to calculate the GCR rate by the method of the median (106).

### PCR and sequencing of GCR survivors

Genomic DNA from GCR survivors was isolated using a Wizard Genomic DNA Purification Kit (Promega). 1 µl of genomic DNA (∼100 ng) was mixed with 8 µl of 1x Cutsmart buffer (New England Biolabs (NEB), Ipswich, MA) and boiled for 10 min at 94ᵒC. 1 µl of tailing mix (0.05 µl Terminal Transferase (NEB, cat. no. M0315), 0.1 µl 10x Cutsmart buffer, 0.1 µl 10 mM dCTP, 0.75 µl dH_2_O) was added and incubated for 30 min at 37ᵒC, 10 min at 65ᵒC, and 5 min at 96ᵒC. Immediately after tailing, 30µl of PCR mix was added. The PCR mix consisted of 4 µl 10x PCR buffer (670 mM Tris-HCl pH 8.8, 160 mM (NH_4_)_2_SO_4_, 50% glycerol, 0.1% Tween-20), 0.32 µl 25 mM dNTP mix, 0.3 µl 100 µM site-specific primer (5’-TTTTCGCCTCGACATCATCTGC-3’), 0.3 µL 100 µM G_18_ primer (5’-CGGGATCCG_18_-3’), 0.5 µl Q5 High-Fidelity DNA Polymerase (NEB, cat. no. M0491), 24.68 µl dH_2_O. The samples were denatured at 98°C for 3 min, followed by 35 cycles of 98°C for 30 s, 68°C for 15 s, and a final extension step at 72°C for 2 min. PCR products were separated on 2.5% agarose gels and extracted using a NucleoSpin Gel and PCR Clean-up kit (Macherey-Nagel, Düren, Germany, cat. no. 740609). The purified PCR products were then cloned using a Zero Blunt TOPO PCR Cloning Kit (Thermo Fisher Scientific, cat. no. 450245). Individual clones were sequenced by Eurofins Genomics (Germany) and the resulting data were analyzed using Sequencher software (Gene Codes, Ann Arbor, MI).

### Genome-wide screen for mutants with increased ITS-GCR rate

The high-throughput replica-pinning screen was performed essentially as previously described (36) using a ROTOR-HDA pinning robot (Singer Instruments). In brief, *prb1ΔhphMX-50bp_ITS* and *hxt13ΔURA3* were introduced into the YKO and ts libraries, with each strain of the libraries present in quadruplicate, by the SGA method (39). The screen was performed at 30°C and 22°C for the YKO and ts strains, respectively. Strains in which less than two colonies (out of a possible four) grew on the final SGA plates were excluded from further analysis. Each plate of the resulting *prb1ΔhphMX-50bp_ITS hxt13ΔURA3 xxx::kanMX* strains (where *xxx::kanMX* indicates either a gene knockout or ts allele) was then replica pinned twice in succession onto six SD-arginine+canavanine+5-FOA plates, yielding 24 colonies per strain. Only hits with a Fisher’s exact test P-value less than 0.002, comparing to the observed number of colonies on selective and non-selective plates for all strains in the screen, were selected for validation, which occurred in two steps. First, for only the screen hits, the *prb1ΔhphMX-50bp_ITS hxt13ΔURA3 xxx::kanMX* strains were generated again by the SGA method. Each strain was then streaked in a 1 cm x 1.5 cm patch on an SD-uracil plate, incubated at 30°C (for YKO strains) or 22°C (for ts strains) for 24-48 h, replica-plated on SD-arginine+canavanine+5-FOA to detect GCR events, and scored by visual inspection. Twenty-one strains (5 YKO and 16 ts) passed this validation step. These YKO or ts mutants were tested with one fluctuation test, and if an increase in ITS-GCR rate was observed, the mutants were introduced into a different strain background (W303), and then subsequently subjected to at least three additional fluctuation tests.

### Rap1 immunoblotting

Cells were harvested and fixed using 20% trichloroacetic acid (TCA). A FastPrep 5G system (MP Biomedicals) was employed for cell disruption, following the recommended program for *S. cerevisiae*. An additional 5% TCA was added to adjust the final TCA concentration to 7.3%. The samples were then centrifuged at 14,000 rpm for 10 min at 4°C, after which the supernatant was discarded. Samples were resuspended in 1x loading buffer and loaded on 10% polyacrylamide gels. Proteins were transferred onto polyvinylidene difluoride membranes. The membranes were blocked in 2.5% bovine serum albumin (BSA) for 60 min at room temperature and incubated with Rap1 antibody (sc-374297; Santa Cruz Biotechnology) in 2.5% BSA overnight at 4°C. The membranes were washed three times with 1× TBS containing 0.1% Tween 20 (Sigma) and incubated with m-IgGκ BP-HRP antibody (sc-516102; Santa Cruz Biotechnology) in 2.5% BSA for 2 h at room temperature. Blots were visualized using the Gel Doc XR+ System (Bio-Rad Laboratories).

### Liquid culture senescence assay

Liquid culture senescence assays were performed essentially as previously described (107). Each senescence assay started with diploid strains. Freshly dissected haploid spores were allowed to form colonies on YPD agar plates after two days of growth at 30°C or 22°C. Cells from these colonies were serially passaged in liquid culture medium at 24-h intervals. For each passage, the cell density of each culture was measured by optical density (calibrated by cell counting using a haemocytometer), and the cultures were diluted back into fresh medium at a cell density of 2 × 10^5^ cells/ml. Cell density was plotted as a function of population doublings.

### Genome-wide screen for mutants with decreased ITS-GCR rate

To screen for mutants with decreased ITS-GCR rate, the high-throughput replica-pinning screen was repeated with a few modifications. In brief, the SGA-derived *prb1ΔhphMX-50bp_ITS hxt13ΔURA3 xxx::kanMX* strains were replica pinned onto six YPD+G418+hygromycin B, yielding 24 colonies per strain. The strains were then re-pinned twice in succession onto SD-arginine+canavanine+5-FOA plates. Only hits with a Fisher’s exact test P-value less than 10^-4^ were selected for validation. A lower P-value was used due to the higher number of hits. Validation occurred in several steps. First, for only the screen hits, the *prb1ΔhphMX-50bp_ITS hxt13ΔURA3 xxx::kanMX* strains were generated again by the SGA method. Each strain was then streaked in a 1 cm x 1.5 cm patch on an SD-uracil plate, incubated at 30°C (for YKO strains) or 22°C (for ts strains) for 24-48 h, replica-plated on SD-arginine+canavanine+5-FOA to detect GCR events, and scored by visual inspection. Second, the strains were generated again by the SGA method, but with an extra copy of *CIN8* (*ho::CIN8-natMX*) also introduced into the YKO and ts strains. These strains were also tested using the patch-and-replica-plate approach. Third, a GCR survivor (ZYY261) was generating by growing FRY910 (the SGA starting strain with the 50-bp ITS and GCR markers) on media containing canavanine and 5-FOA. ZYY261 was confirmed by PCR and sequencing to have a de novo telomere added at the ITS site. ZYY261 was crossed to the hits from the screen using the SGA method, and resulting strains that showed a synthetic genetic growth defect by visual comparison of colony sizes were eliminated from further analysis. Lastly, due to the large number of hits that passed the first three validation steps, only a subset of these was subsequently subjected to at least three fluctuation tests, and the mutants were not tested in a different strain background.

### Gene ontology enrichment analysis

The GO term finder tool (http://go.princeton.edu/) was used to query biological process enrichment for each gene set, with a P-value cutoff of 0.01 and Bonferroni correction applied. REVIGO (108) was used to further analyze the GO term enrichment data, using the “Medium (0.7)” term similarity filter and the simRel score as semantic similarity measure. As a result, terms with a frequency more than 10% in the REVIGO output were eliminated for being too broad.

## Acknowledgments and funding sources

We thank G.W. Brown for providing yeast strains; D. Novarina and L.M. Veenhoff for critical reading of the manuscript. F.R.R.B. was supported by a Consejo Nacional de Ciencia y Tecnología (CONACYT) scholarship. Z. Yin was supported by a scholarship from the Nanjing Huimou Medi-Tech Co. Y.Y. was supported by scholarships from the University of Groningen’s Abel Tasman Talent and from the Chongqing Explorer Decoration Engineering Co. Work in the laboratory of M. Chang was supported by an Open Competition M-2 grant from the Dutch Research Council.

---

**Table S1.**
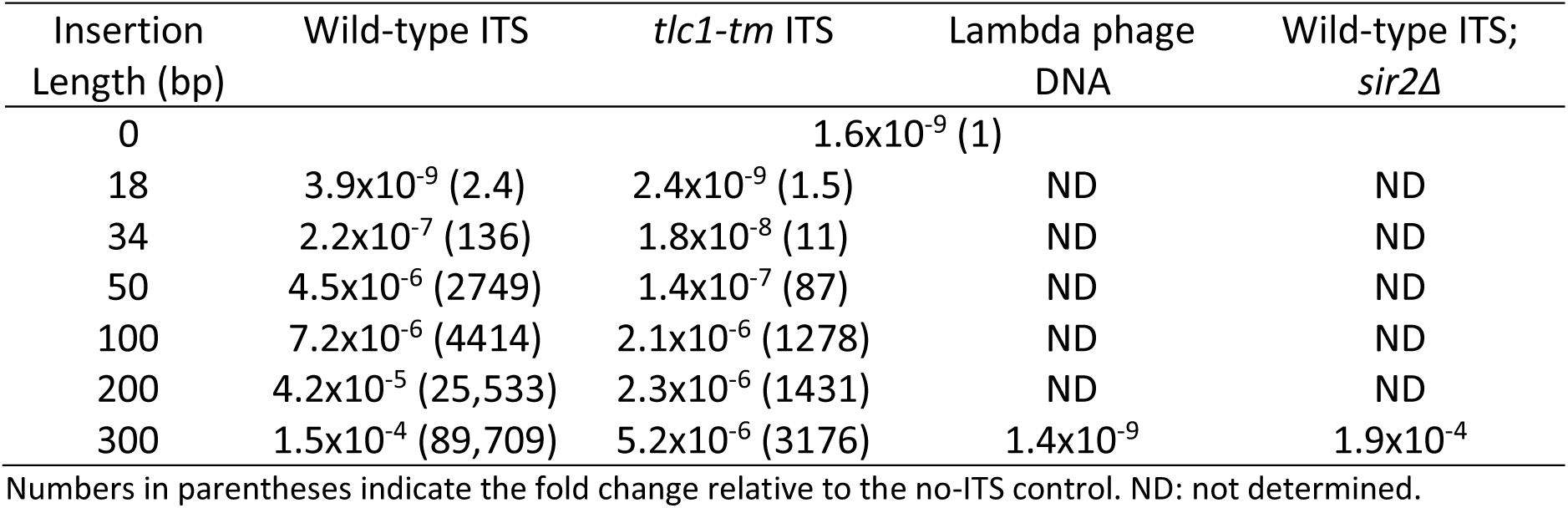
GCR rate increases with ITS length.

**Table S2.**
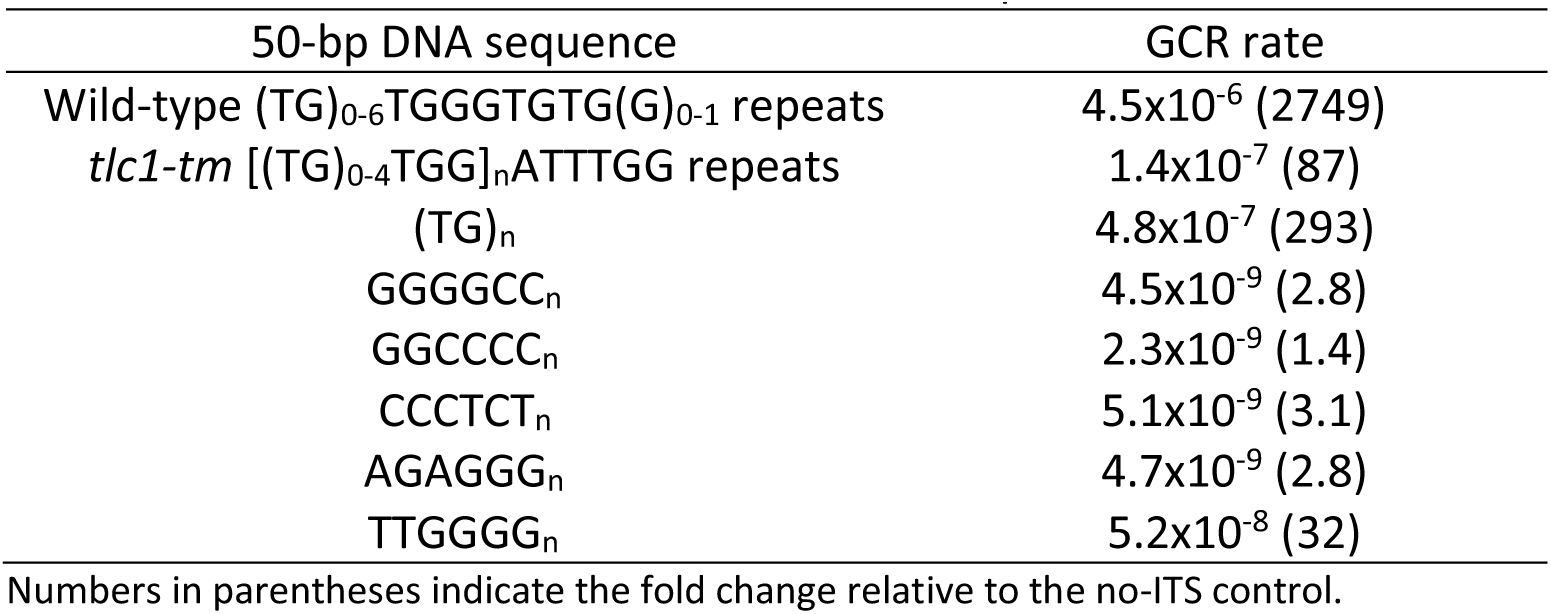
GCR rates of strains with different 50-bp insertions.

**Table S3.** GCR rates of mutants identified in the ITS-GCR screen without and with a 50-bp ITS.

| Genotype | No ITS | 50-bp ITS |
| --- | --- | --- |
| 30°C |  |  |
| Wild type | 3.5x10<sup>-10</sup> (1)<sup>a</sup> | 4.5x10<sup>-6</sup> (1) |
| *elg1Δ* | 1.6x10<sup>-7</sup> (45)<sup>a</sup> | 1.4x10<sup>-5</sup> (3.0) |
| *rad27Δ* | 3.6x10<sup>-7</sup> (1040)<sup>a</sup> | 1.6x10<sup>-4</sup> (36) |
| *ydl162cΔ* | 4.6x10<sup>-9</sup> (13)<sup>a</sup> | 3.4x10<sup>-5</sup> (7.6) |
| 22°C |  |  |
| Wild type | 1.9x10<sup>-9</sup> (1)<sup>b</sup> | 2.3x10<sup>-6</sup> (1) |
| *cdc2-2* | 1.4x10<sup>-8</sup> (7.5)<sup>b</sup> | 1.1x10<sup>-5</sup> (4.7) |
| *cdc2-7* | 2.3x10<sup>-8</sup> (12)<sup>b</sup> | 1.2x10<sup>-5</sup> (5.3) |
| *cdc9-1* | 5.4x10<sup>-7</sup> (283) | 1.2x10<sup>-5</sup> (5.5) |
| *rfc2-1* | 6.8x10<sup>-9</sup> (3.5)<sup>b</sup> | 7.4x10<sup>-6</sup> (3.3) |
| *rfc4-20* | 4.0x10<sup>-7</sup> (209)<sup>b</sup> | 2.1x10<sup>-5</sup> (9.3) |
| *rfc5-1* | 8.7x10<sup>-9</sup> (4.5)<sup>b</sup> | 5.9x10<sup>-6</sup> (2.6) |
| *rpt4-150* | 1.6x10<sup>-9</sup> (0.9)<sup>b</sup> | 1.2x10<sup>-5</sup> (5.5) |

Numbers in parentheses indicate the fold change relative to the corresponding wild-type control.
<sup>a</sup>Data taken from a previous publication using the original GCR assay without an ITS (1).
<sup>b</sup>These values were derived from strains without an ITS, but with the *hphMX* marker inserted at the *PRB1* locus.

**Table S4.** GCR rates of selected mutants not identified in the ITS-GCR screen without and with a 50-bp ITS.

| Genotype | No ITS<sup>a</sup> | 50-bp ITS |
| --- | --- | --- |
| Wild type | 3.5x10<sup>-10</sup> (1) | 4.5x10<sup>-6</sup> (1) |
| *pif1Δ* | 3.5x10<sup>-7</sup> (1010) | 1.1x10<sup>-5</sup> (2.5) |
| *rad50Δ* | 2.3x10<sup>-7</sup> (657) | 3.3x10<sup>-6</sup> (0.72) |
| *sic1Δ* | 1.8x10<sup>-7</sup> (514) | 4.5x10<sup>-6</sup> (1.0) |
| *rlf2Δ* | 1.2x10<sup>-7</sup> (343) | 4.0x10<sup>-6</sup> (0.9) |
| *rmi1Δ* | 6.0x10<sup>-8</sup> (189) | 7.4x10<sup>-6</sup> (1.6) |
| *mus81Δ* | 6.5x10<sup>-8</sup> (186) | 4.5x10<sup>-6</sup> (1.0) |
| *rad52Δ* | 4.0x10<sup>-8</sup> (115) | 5.4x10<sup>-5</sup> (12.1) |
| *slx8Δ* | 2.6x10<sup>-8</sup> (75) | 6.8x10<sup>-6</sup> (1.5) |
| *slx5Δ* | 2.3x10<sup>-8</sup> (66) | 1.2x10<sup>-5</sup> (2.7) |
| *sgs1Δ* | 1.2x10<sup>-8</sup> (34) | 3.7x10<sup>-6</sup> (0.81) |
| *rrm3Δ* | 1.4x10<sup>-9</sup> (4.0) | 1.7x10<sup>-5</sup> (3.8) |

Numbers in parentheses indicate the fold change relative to the wild-type control.
<sup>a</sup>Data taken from a previous publication using the original GCR assay without an ITS (1).

**Table S5.**
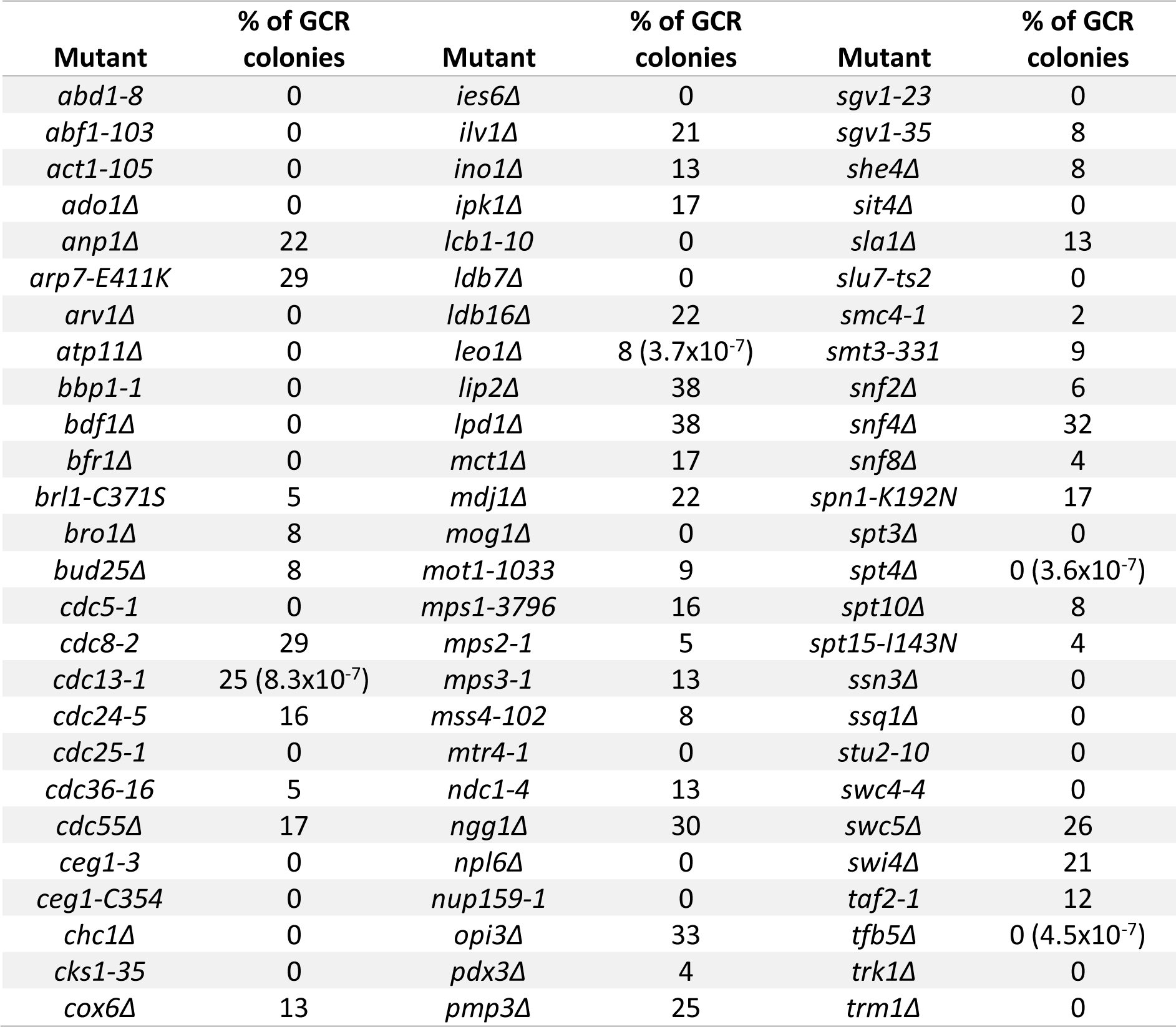
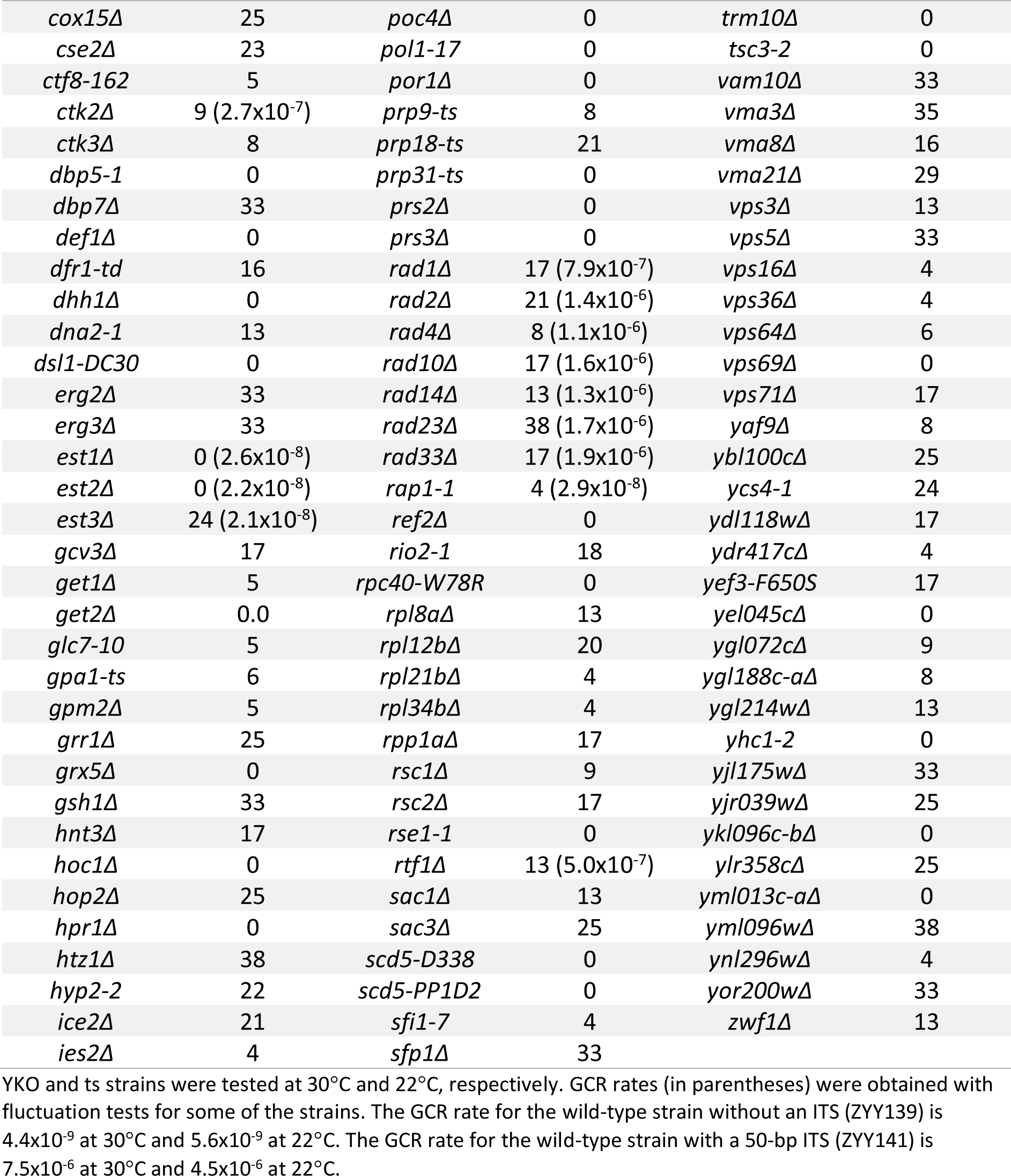
List of mutants identified in the screen that decrease ITS-induced GCRs.

**Table S6.** GCR rates of telomere and NER mutants with a 50-bp ITS.

| Genotype | GCR rate |
| --- | --- |
| 30°C |  |
| Wild type without ITS | 4.4x10<sup>-9</sup> (1) |
| Wild type | 7.5x10<sup>-6</sup> (1720) |
| *est1Δ* | 2.5x10<sup>-8</sup> (5.8) |
| *est2Δ* | 2.2x10<sup>-8</sup> (5.0) |
| *est3Δ* | 2.1x10<sup>-8</sup> (4.8) |
| *rad4Δ* | 1.1x10<sup>-6</sup> (262) |
| *rad23Δ* | 1.7x10<sup>-6</sup> (389) |
| *rad33Δ* | 1.9x10<sup>-6</sup> (428) |
| *rad14Δ* | 1.3x10<sup>-6</sup> (301) |
| *rad1Δ* | 9.8x10<sup>-7</sup> (225) |
| *rad10Δ* | 1.6x10<sup>-6</sup> (362) |
| *rad2Δ* | 1.4x10<sup>-6</sup> (315) |
| *tfb5Δ* | 4.5x10<sup>-7</sup> (103) |
| ***rad7Δ*** | 6.5x10<sup>-6</sup> (1493) |
| ***rad16Δ*** | 6.2x10<sup>-6</sup> (1419) |
| ***rad26Δ*** | 2.3x10<sup>-6</sup> (530) |
| 22°C |  |
| Wild type without ITS | 5.6x10<sup>-9</sup> (1) |
| Wild type | 4.5x10<sup>-6</sup> (815) |
| *cdc13-1* | 8.3x10<sup>-7</sup> (148) |
| *rap1-1* | 2.9x10<sup>-8</sup> (5.1) |

Numbers in parentheses indicate the fold change relative to the no-ITS control.
Gene deletions in bold were not identified in the screen.

**Table S7.**
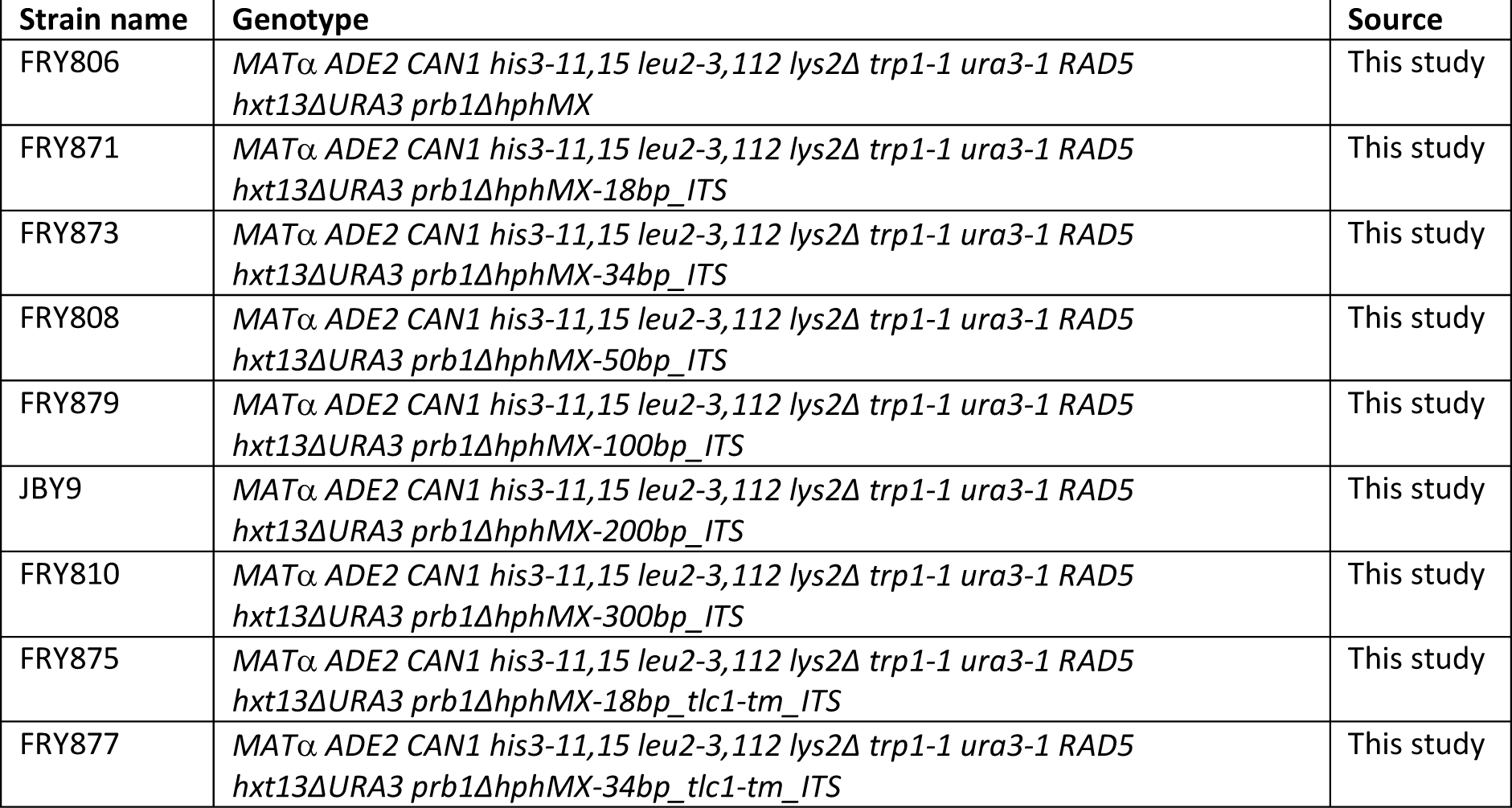
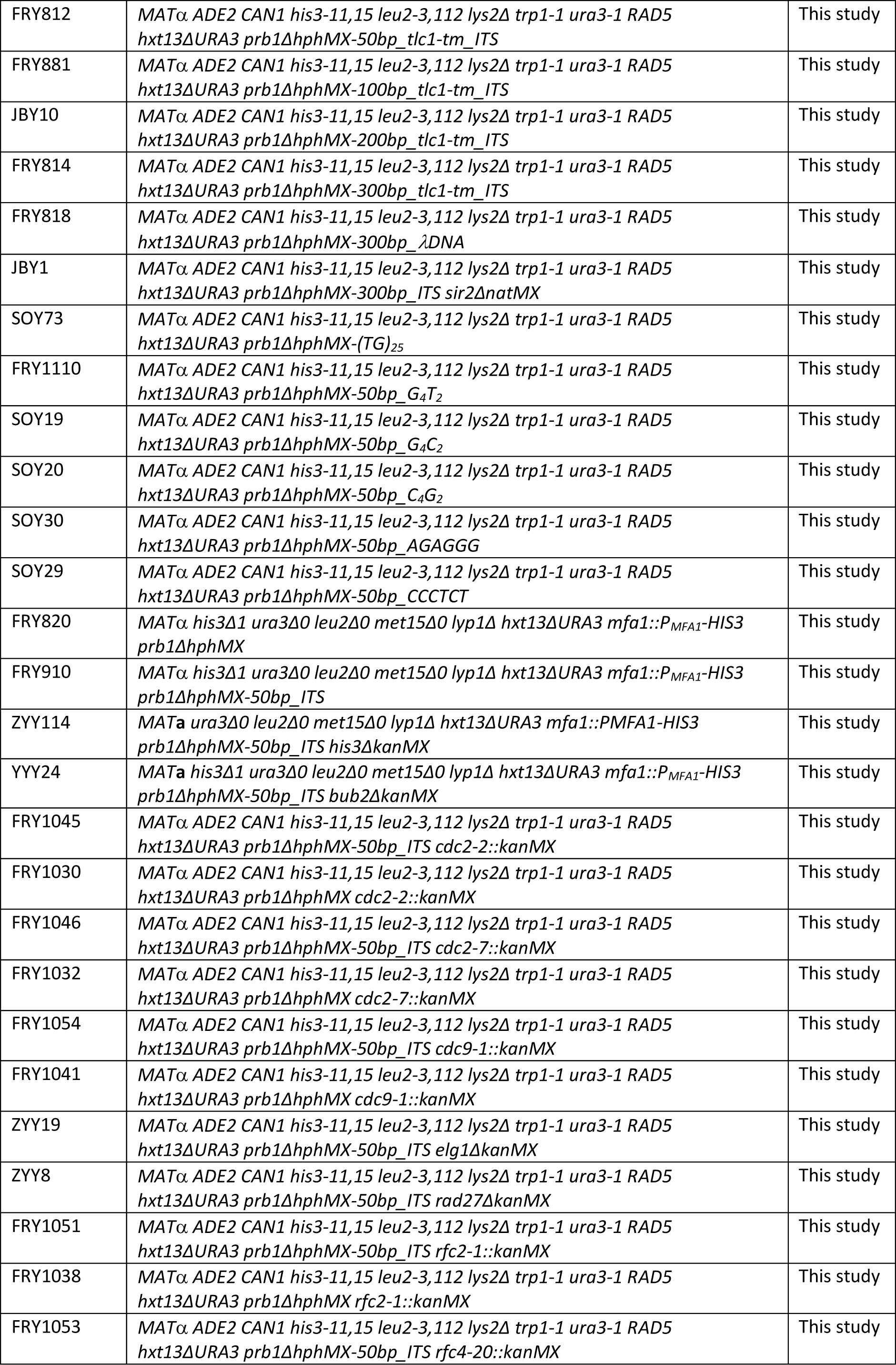
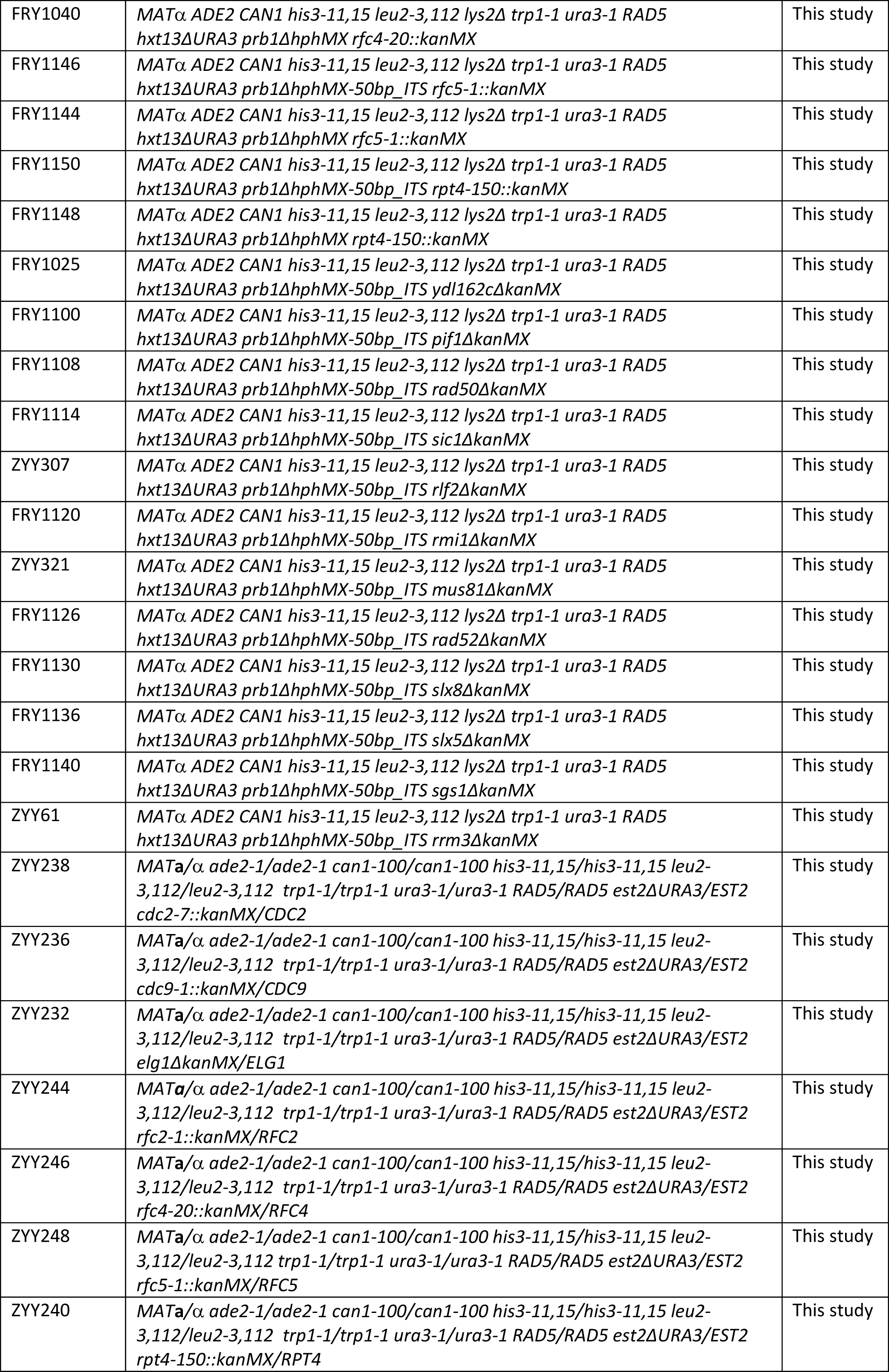
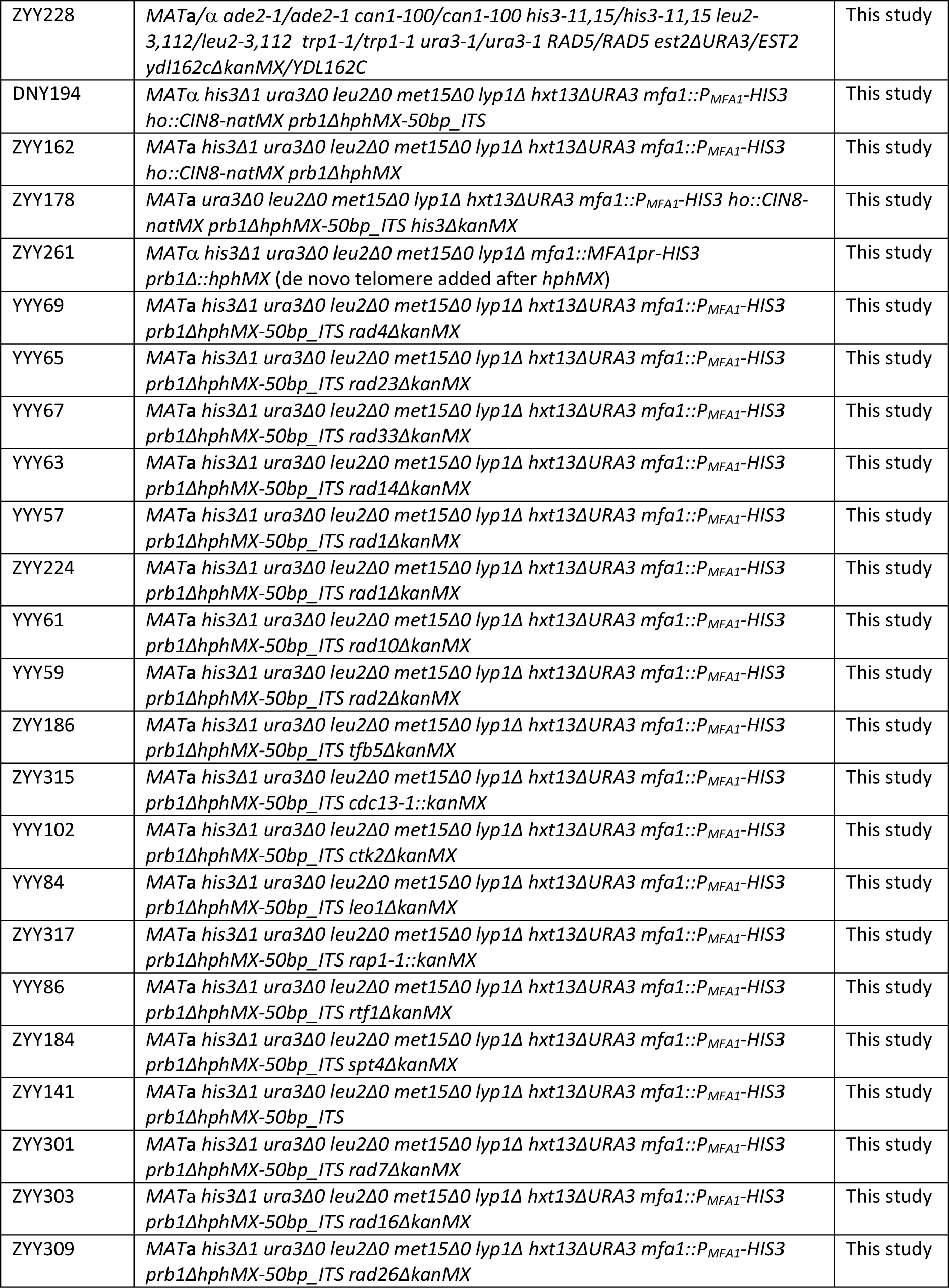
Yeast strains used in this study.

**Table S8.**
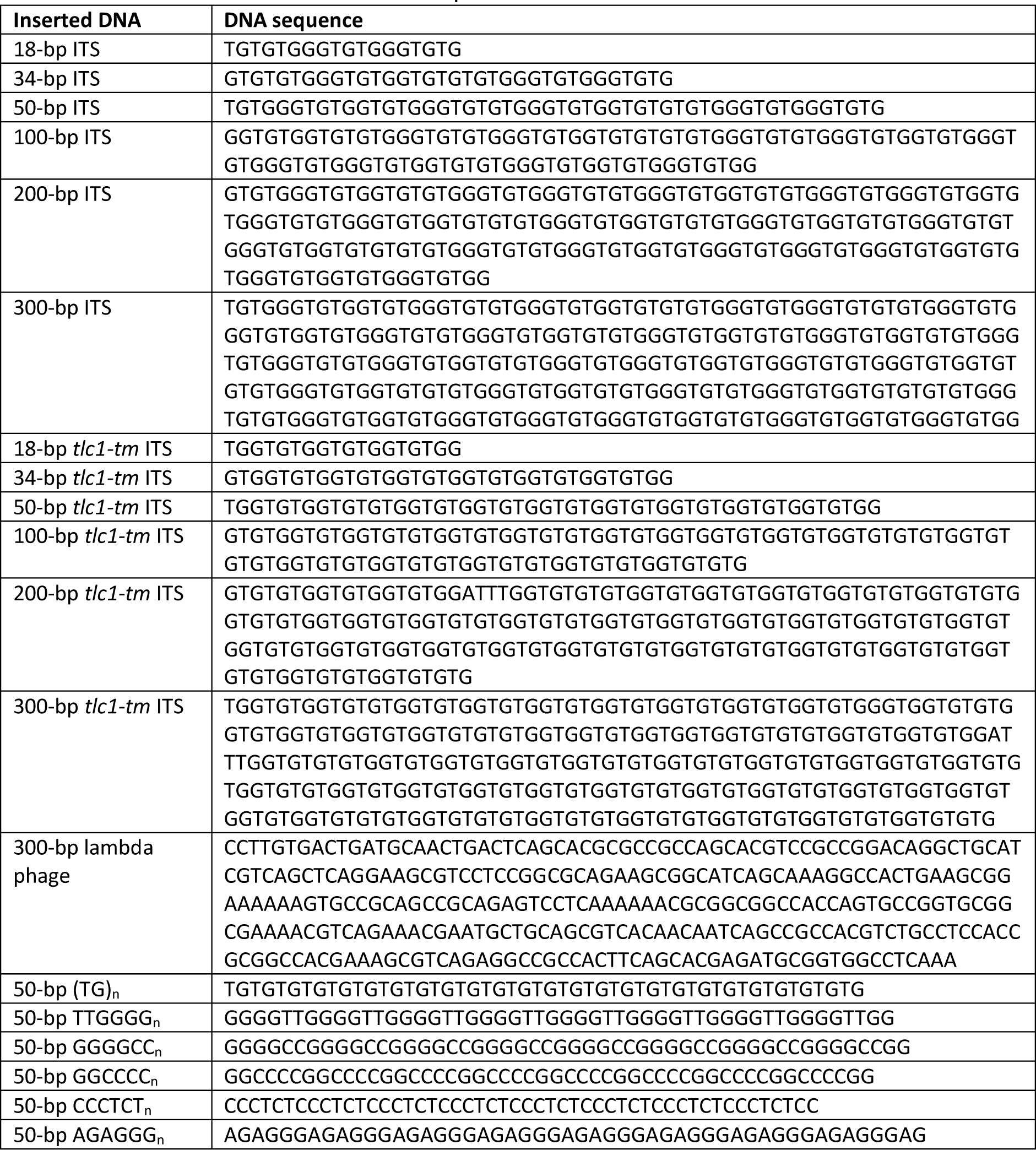
ITS and other inserted DNA sequences.

**Figure S1.**
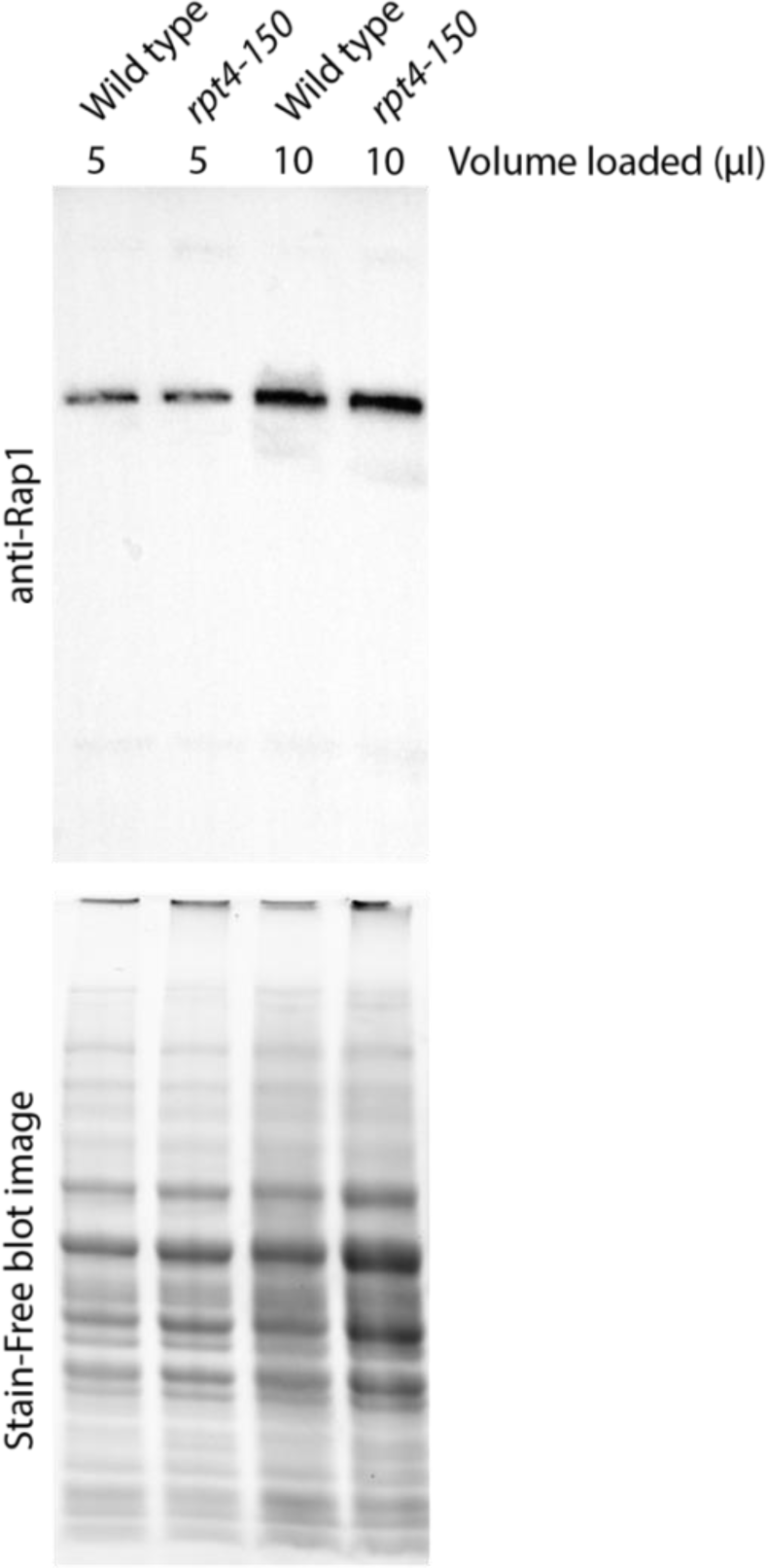
Rap1 protein levels in wild-type and *rpt4-150* strains. Wild-type and *rpt4-150* cells, harvested from logarithmically growing cultures, were fixed using TCA. Extracts from each sample were subjected to SDS-PAGE separation, followed by immunoblotting to detect Rap1 protein. Stain-Free imaging was used to assess the total protein input.

**Figure S2.**
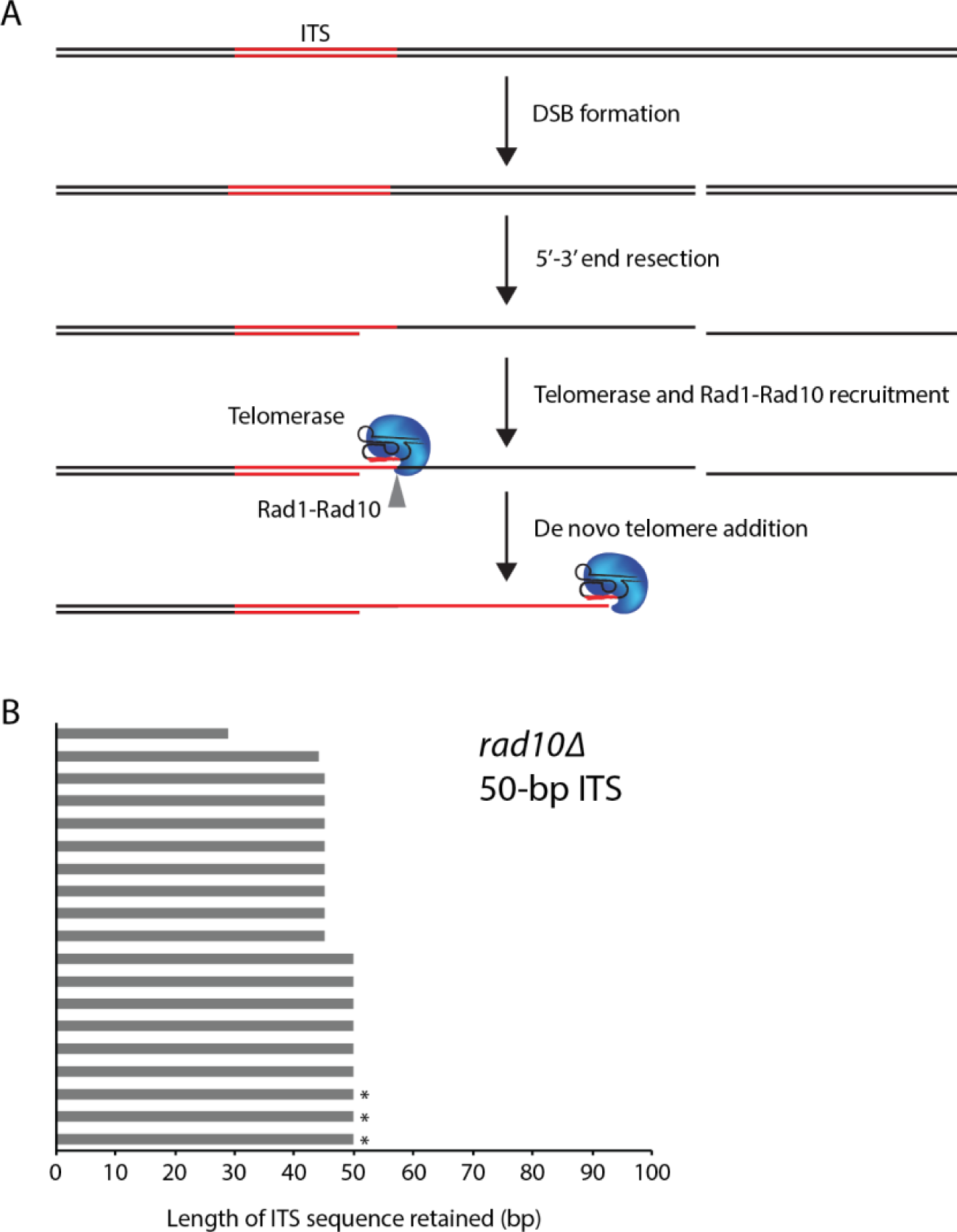
Deletion of *RAD10* does not eliminate GCR survivors that retain the entire ITS. (**A**) Model for how the Rad1-Rad10 endonuclease could function during de novo telomere addition. A break that occurs distal to the ITS will undergo 5’–3’ end resection until a portion of the ITS becomes single stranded, which will then bind Cdc13 (not shown). Cdc13 recruits telomerase, followed by Rad1-Rad10 cleavage of the 3’ overhang/flap, allowing telomerase to extend the ITS into a de novo telomere. Deletion of *RAD1* or *RAD10* should reduce or eliminate GCR survivors that retain the entire ITS in the de novo telomere. (**B**) GCR survivors were isolated by growing *rad10Δ* strain with a 50-bp ITS on agar plates containing canavanine and 5-FOA. Nineteen independent isolates were analyzed. In all isolates, the presence of a de novo telomere added at the ITS site was confirmed by PCR and sequencing. The length of the original ITS retained for each de novo telomere is plotted. Nine out of 19 retain the entire ITS, including three isolates that also retained up to 14 bp of sequence downstream of the ITS (indicated by asterisks), contrary to what the model predicts.

**Figure S3.**
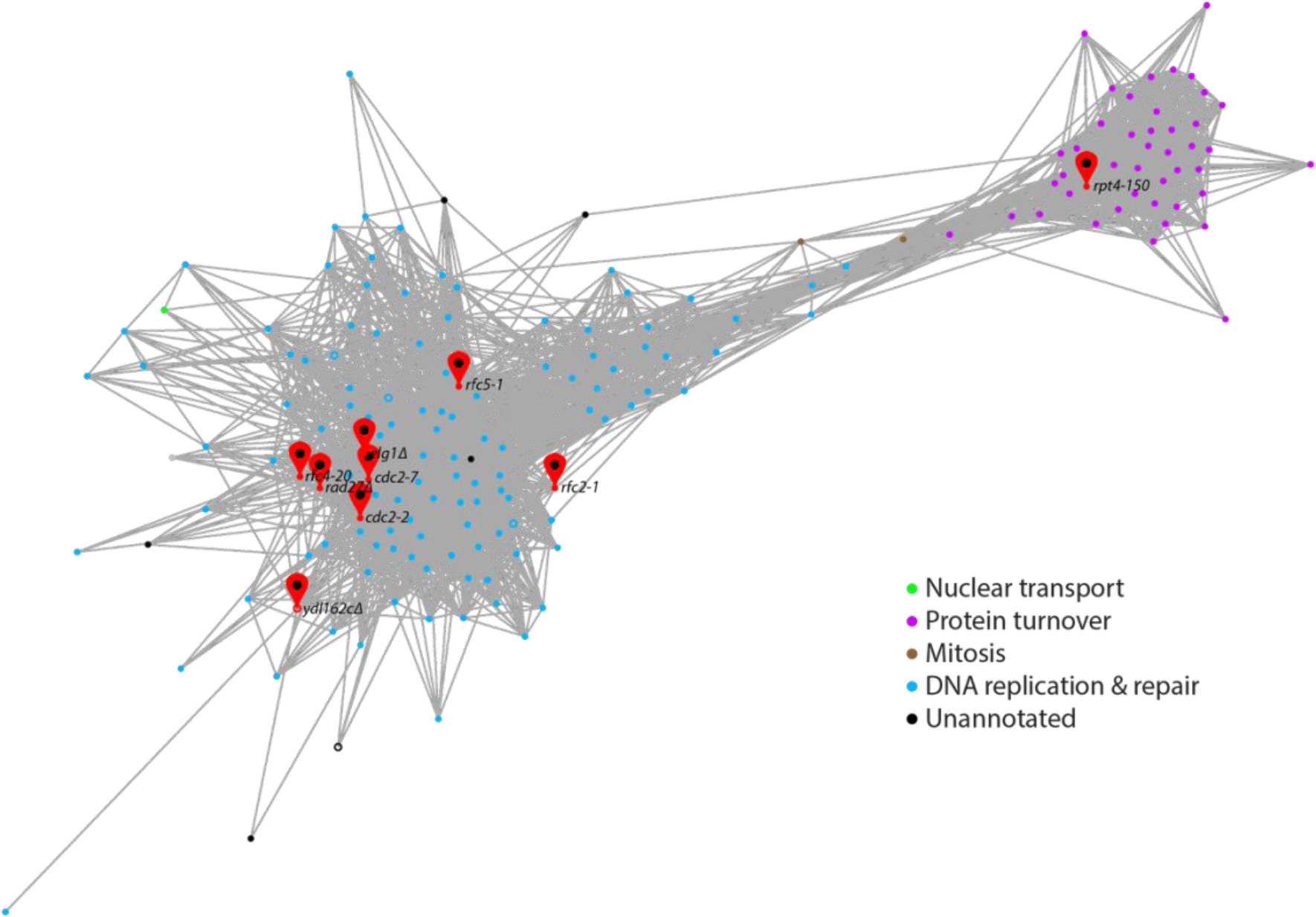
Genetic interaction profile similarity subnetwork for genes that suppress ITS-induced GCR rate. The network was generated using TheCellMap.org (2). Nodes (representing deletions of nonessential genes or temperature-sensitive alleles of essential genes) sharing similar genetic interaction profiles (PCC > 0.2) are connected by an edge in the network. Genes sharing similar genetic interaction profiles map closer to each other. The subnetwork was annotated using Spatial Analysis of Functional Enrichment (SAFE; 3).

